# MEILB2-BRME1 forms a V-shaped DNA clamp upon BRCA2-binding in meiotic recombination

**DOI:** 10.1101/2023.07.04.547728

**Authors:** Manickam Gurusaran, Jingjing Zhang, Kexin Zhang, Hiroki Shibuya, Owen R. Davies

## Abstract

DNA double-strand break repair by homologous recombination has a specialised role in meiosis by generating crossovers that enable the formation of haploid germ cells. This requires meiosis-specific MEILB2-BRME1, which interacts with BRCA2 to facilitate loading of recombinases onto resected DNA ends. Here, we report the crystal structure of the MEILB2-BRME1 2:2 core complex, revealing a parallel four-helical assembly that recruits BRME1 to meiotic double-strand breaks *in vivo*. It forms an N-terminal β-cap that binds to DNA, and a MEILB2 coiled-coil that bridges to C-terminal ARM domains. Upon BRCA2-binding, MEILB2-BRME1 2:2 complexes dimerize into a V-shaped 2:4:4 complex, with rod-like MEILB2-BRME1 components arranged at right-angles. The β-caps located at the tips of the MEILB2-BRME1 limbs are separated by 25 nm, allowing them to bridge between DNA molecules. Thus, we propose that BRCA2 induces MEILB2-BRME1 to function as a DNA clamp, connecting resected DNA ends or homologous chromosomes to facilitate meiotic recombination.

## Introduction

Breast cancer susceptibility protein BRCA2 performs a central role in recombination-mediated DNA double-strand break (DSBs) repair by loading recombinases onto resected DNA ends (Jensen et al., 2010, Liu et al., 2010, Thorslund et al., 2010). Its importance in genome stability is demonstrated by the gross chromosomal rearrangements that accumulate in BRCA2 deficient mammalian cells (Patel et al., 1998, Yu et al., 2000), and the strong association between germline BRCA2 mutations and early-onset breast and ovarian cancers (Hall et al., 1990). BRCA2 is also required for meiosis, as its germline-specific deficiency leads to meiotic impairment and infertility in mice (Sharan et al., 2004). Here, programmed DSBs are induced, and then repaired by recombination between homologous chromosomes to generate crossovers, which provide genetic diversity and are critical for the correct segregation of homologues (Hunter, 2015, Zickler and Kleckner, 2015). Hence, BRCA2 is essential for maintaining genome integrity in somatic cells and for generating haploid germ cells in meiosis.

In somatic cells, homologous recombination is initiated by DSB end resection, in which the MRN complex generates 5’ single-stranded DNA (ssDNA) overhangs that become coated by RPA (Gnugge and Symington, 2021) (Figure 1). BRCA2 then displaces RPA by loading recombinase RAD51 to form RAD51-ssDNA filaments (Shahid et al., 2014, Bell et al., 2023). These nucleoprotein filaments invade the sister chromatid and mediate template-directed repair, generating joint molecules that are resolved by multiple pathways (Kowalczykowski, 2015). In meiosis, homologous recombination uses the same principal machinery, with several critical modifications (Figure 1): 1. Meiotic DSBs are induced by the SPO11-TOPOVIBL complex (Keeney et al., 1997, Robert et al., 2016). 2. Resected ssDNA overhangs are coated by RPA and meiosis-specific complex MEIOB-SPATA22 (Souquet et al., 2013, Ribeiro et al., 2021, Luo et al., 2013). 3. BRCA2 loads RAD51 and the meiosis-specific recombinase DMC1 onto ssDNA overhangs (Pittman et al., 1998, Dai et al., 2017, Shinohara and Shinohara, 2004). 4. BRCA2 and recombinases are supported by essential meiosis-specific accessory proteins, including MEILB2-BRME1 (Zhang et al., 2022). 5. Meiotic recombination uses the homologous chromosome rather than the sister chromatid as the primary repair template (Lao and Hunter, 2010). 6. Meiotic recombination occurs within the synaptonemal complex that binds together homologous chromosomes (Adams and Davies, 2023). 7. The processing of recombination intermediates is tightly regulated to ensure the formation of at least one crossover, and rarely more than two crossovers, per homologous chromosome pair (Hunter, 2015, Zickler and Kleckner, 2015).

**Figure 1.**
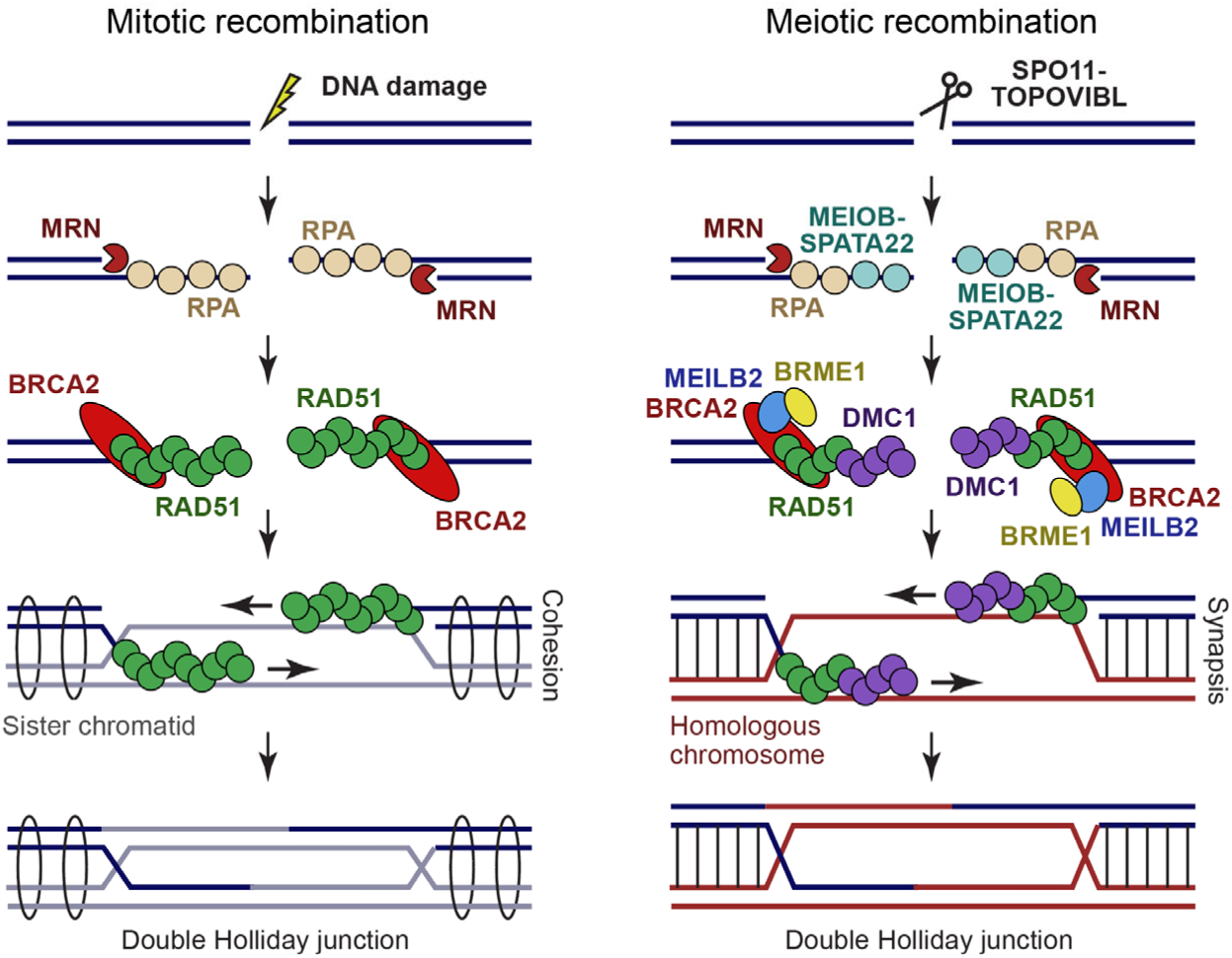
Mechanics of homologous recombination. Schematic of the key molecular players in mitotic (left) and meiotic (right) recombination. The main specialisations of meiotic recombination are DSB induction by SPO11-TOPOVIBL, coating of resected ends with MEIOB-SPATA22 (alongside RPA), the requirement for recombinase DMC1 (in addition to RAD51), and the role of MEILB2-BRME1 in BRCA2 function. These facilitate use of the synapsed homologous chromosome, rather than the cohesin-bound sister chromatid, as the primary repair template. In meiotic recombination, double Holliday junctions are resolved into crossovers and non-crossovers in a highly regulated manner. In mitotic recombination, crossovers are less frequent, and additional pathways operate that bypass the double Holliday junction intermediate.

BRCA2 is a large protein (3,329 amino-acids in mice) that contains multiple recombinase-binding sites and a single globular DNA-binding domain (Figure 2a). BRC repeats within the central exon 11 region bind to RAD51 and DMC1 by competing with their oligomerisation (Pellegrini et al., 2002, Martinez et al., 2016). In contrast, sequences that flank the DNA-binding domain (exons 14 and 27), interact with DMC1 and RAD51, respectively, in a manner that stabilises their nucleoprotein filaments (Davies and Pellegrini, 2007, Esashi et al., 2007, Thorslund et al., 2007). Finally, the DNA-binding domain contains OB-folds for ssDNA-binding (Yang et al., 2002), and interacts with acidic DNA-mimicking protein DSS1 (Marston et al., 1999), which act together to offload RPA from ssDNA overhangs (Zhao et al., 2015, Bell et al., 2023). Hence, whilst we lack a full molecular mechanism, BRCA2 contains the necessary functionalities to explain its role in displacing RPA, and loading recombinases onto ssDNA overhangs to form nucleoprotein filaments (Jensen et al., 2010, Liu et al., 2010, Thorslund et al., 2010).

**Figure 2.**
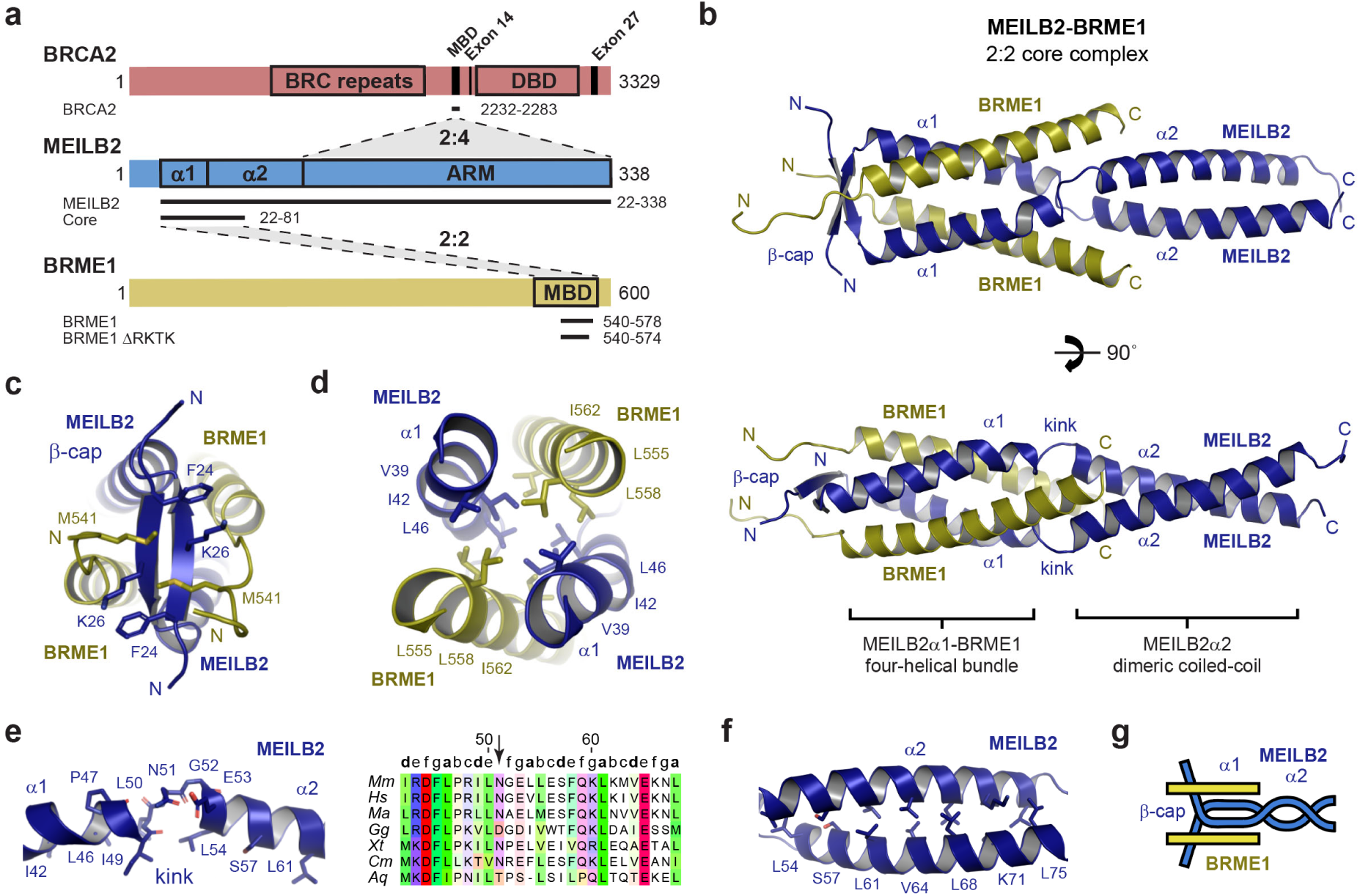
Crystal structure of the MEILB2-BRME1 2:2 core complex. **(a)** Schematic of mouse BRCA2, MEILB2 and BRME1 sequences, highlighting domains, interaction sites and constructs used in this study. BRCA2 includes recombinase-binding BRC repeats, a MEILB2-binding domain (MBD), DMC1-binding exon 14, a DNA-binding domain (DBD) and RAD51-binding exon 27. MEILB2 contains a BRME1-binding α-helical region, and BRCA2-binding armadillo repeat (ARM) domain. BRME1 has a MEILB2-binding domain (MBD). (**b**) Crystal structure of the MEILB2-BRME1 2:2 core complex. MEILB2 α1 (blue) and BRME1 (yellow) α-helices interact within a parallel four-helical bundle. MEILB2 α1 helices are followed by kinks that re-orientate α2 helices to form a parallel dimeric coiled-coil, and are preceded by a β-cap. (**c**) The N-terminal β-cap is a two-stranded anti-parallel β-sheet that is oriented perpendicular to the coiled-coil axis, supported by hydrophobic interactions of MEILB2 F24 and K26, and BRME1 M541. (**d**) The hydrophobic core of the parallel four-helical bundle is formed of leucine, isoleucine and valine amino-acids of MEILB2 and BRME1 chains, as indicated. (**e**) The kink within the MEILB2 chain is defined by non-helical backbone torsion angles of L50 and N51 (left), and a single amino-acid insertion between heptad repeats of α1 and α2 helices (indicated by an arrow, right). Multiple sequence alignment: *Mus musculus* (*Mm*), *Homo sapiens* (*Hs*), *Mesocricetus auratus* (*Ma*), *Gallus gallus* (*Gg*), *Xenopus tropicalis* (*Xt*), *Callorhinchus milii* (*Cm*), and *Amphimedon queenslandica* (*Aq*). (**f**) The MEILB2 α2 parallel dimeric coiled-coil is formed of ‘a’ and ‘d’ heptad amino-acids L54, S57, L61, V64, L68, K71 and L75. (**g**) Schematic of the MEILB2-BRME1 2:2 core structure.

MEILB2 (Meiotic localizer of BRCA2; also known as HSF2BP) was recently identified as a meiosis-specific interacting partner of BRCA2 (Zhang et al., 2019, Brandsma et al., 2019). BRME1 (BRCA2 and MEILB2-associating protein 1; also known as MEIOK21, MMER, C19ORF57 and 2930432K21Rik) was then identified as a meiosis-specific binding partner of MEILB2 (Zhang et al., 2020, Felipe-Medina et al., 2020, Takemoto et al., 2020, Shang et al., 2020, Li et al., 2020). MEILB2 and BRME2 co-localise on meiotic recombination sites in a mutually dependent manner (Zhang et al., 2019, Zhang et al., 2020). In MEILB2 and BRME1 knockout mice, males are sterile, with failure of RAD51 and DMC1 foci formation, failure of DSB repair and failure of crossover formation (Zhang et al., 2019, Brandsma et al., 2019, Zhang et al., 2020, Felipe-Medina et al., 2020, Takemoto et al., 2020, Shang et al., 2020, Li et al., 2020). In contrast, females exhibit subfertility, in which the development and number of oocytes is substantially reduced (Zhang et al., 2019, Brandsma et al., 2019, Zhang et al., 2020, Felipe-Medina et al., 2020, Takemoto et al., 2020, Shang et al., 2020, Li et al., 2020, Shang et al., 2022). This resembles the sexual dimorphism observed upon deficiency of BRCA2 and other recombination factors (Sharan et al., 2004). Their co-localisation and similar phenotypes suggest that MEILB2 and BRME1 exist within the same functional unit in meiosis. This likely occurs through modulation of BRCA2 function as MEILB2 or BRME1 expression suppresses BRCA2-mediated recombination in somatic cells (Sato et al., 2020, Zhang et al., 2020), and MEILB2 expression has been observed in cancer cell lines and human tumour samples (Brandsma et al., 2019, Sato et al., 2020). Indeed, it has been hypothesised that MEILB2-BRME1 may module the oligomeric state or conformation of BRCA2 to favour meiotic rather than mitotic recombination (Zhang et al., 2022).

The mechanical basis of recombination is an intrinsically structural problem that will likely be solved by understanding the functional architecture of its principal components and complexes. MEILB2 consists of N-terminal coiled-coil and C-terminal ARM (armadillo repeat) domains, which bind with high affinity to BRME1 and BRCA2, respectively (Zhang et al., 2020, Pendlebury et al., 2021) (Figure 2a). The ARM domains interact with the MBD (MEILB2-binding domain) of BRCA2, which is located immediately upstream of its DMC1-binding site (Zhang et al., 2020) (Figure 2a). Crystal structures have revealed that BRCA2-binding staples together two MEILB2-ARM dimers at approximately 90° to one another (Pendlebury et al., 2021, Ghouil et al., 2021). In these 2:4 complexes, each BRCA2 MBD peptide bridges between interacting MEILB2 ARM dimers, through cryptic repeats within their N-and C-terminal ends binding to opposing ARM domains (Pendlebury et al., 2021, Ghouil et al., 2021). The N-terminus of MEILB2’s coiled-coil binds to the C-terminal end of BRME1 (Zhang et al., 2020), which currently has no other known structure or interacting partners. The MEILB2 coiled-coil exists in dimeric and higher-order species in isolation, but forms stable 2:2 complexes with BRME1 (Zhang et al., 2020). However, the lack of structural information regarding the MEILB2-BRME1 complex currently prevents us from understanding the architecture of the wider BRCA2-MEILB2-BRME1 assembly, and critically the molecular basis of MEILB2-BRME1 function in BRCA2-mediated meiotic recombination.

Here, we report the crystal structure of the MEILB2-BRME1 2:2 core complex, revealing a four-helical assembly that mediates BRME1 recruitment to meiotic DSBs *in vivo*. We show that its N-terminal β-cap binds to DNA, requiring both MEILB2 and BRME1 amino-acids. We report a model of the full MEILB2-BRME1 2:2 complex, and show that BRCA2-binding induces dimerization of this structure into a V-shaped assembly. The β-caps at the ends of the MEILB2-BRME1 limbs of this structure are separated by 25 nm, so bind together distinct DNA molecules, forming protein-DNA networks. Hence BRCA2-binding can induce MEILB2-BRME1 to act as a DNA clamp in meiotic recombination.

## Results

### Crystal structure of MEILB2-BRME1

MEILB2 possesses an N-terminal α-helical region, consisting of two predicted α-helices (α1 and α2), followed by a C-terminal ARM domain (Figure 2a). We previously demonstrated that MEILB2’s α-helical region binds via its α1 helix to the C-terminus of BRME1 (MEILB2-binding domain, MBD), forming a stable 2:2 complex (Zhang et al., 2020). Here, we identified an optimised construct for crystallography, which includes the α1 helix and the beginning of the α2 helix of MEILB2 (amino-acids 22-81; herein referred to as MEILB2 core) and the core of BRME1’s MBD (amino-acids 540-578; herein referred to as BRME1) (Supplementary Figure 1). Crystals of this MEILB2-BRME1 core complex diffracted to a resolution limit of 1.50 Å, enabling structure solution by molecular replacement of ideal helical fragments using *ARCIMBOLDO_LITE* (Caballero et al., 2018) (Table 1 and Supplementary Figure 2). This revealed a 2:2 structure, consisting of an N-terminal parallel four-helical bundle of MEILB2 α1 and BRME1, followed by a C-terminal MEILB2 α2 parallel dimeric coiled-coil (Figure 2b).

**Table 1.**
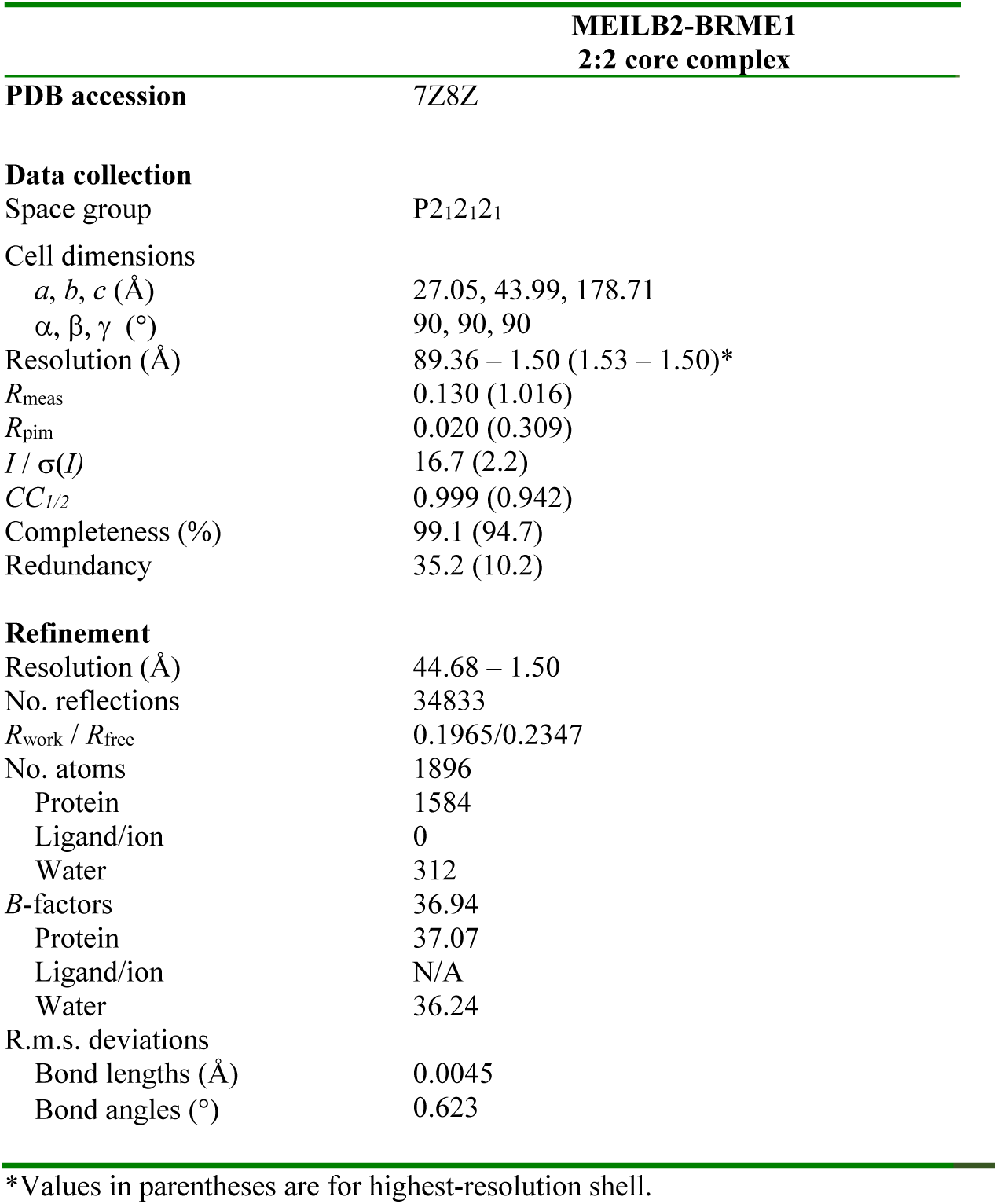
Data collection, phasing and refinement statistics

The MEILB2-BRME1 core structure contains an unusual β-cap at its N-terminal tip (Figure 2b,c). This consists of an anti-parallel two-stranded β-sheet, which is formed by the N-terminal ends of MEILB2 chains, and is orientated perpendicular to the helical axis (Figure 2b,c). The β-cap binds together the N-termini of the two α1 helices of the four-helical bundle, so likely prevents the fraying apart of helical ends that typically occurs within coiled-coil structures (Dragan and Privalov, 2002). Hence, we infer that the β-cap likely confers rigidity to the N-terminal end of the MEILB2-BRME1 2:2 structure. In previous cases of α/β-coiled-coils, the β-sheets typically constitute insertions, as β-layers within the α-helical coiled-coil structure (Hartmann et al., 2016). To our knowledge, MEILB2-BRME1 is the first example of an α/β-coiled-coil in which the β-sheet caps off the end of the α-helical coiled-coil.

Following the β-cap, a MEILB2-BRME1 four-helical bundle is observed, terminating with the splaying apart of the C-termini of BRME1 chains (Figure 2b,d). At this point, there is slight ‘kink’ in the MEILB2 chains, between α1 and α2 helices, where L50 and N51 adopt non-helical conformations due to a single amino-acid insertion into the heptad repeats (Figure 2e). This kink re-orientates the MEILB2 chains from the upstream four-helical bundle to the downstream dimeric coiled-coil conformation (Figure 2b,e). The parallel dimeric coiled-coil of the α2 helix adopts a canonical heptad pattern (Figure 2b,f). Thus, the overall structure of the MEILB2-BRME1 core can be described as a parallel 2:2 α/β-coiled-coil, where an N-terminal MEILB2 β-cap is followed by a MEILB2 α1-BRME1 four-helical bundle that transitions via a kink into a C-terminal MEILB2 α2 dimeric coiled-coil (Figure 2b,g).

### BRME1 recruitment to meiotic DSBs requires its MEILB2-binding interface

To determine whether the observed MEILB2-BRME1 interaction in the crystal structure is crucial for their interaction *in vivo*, we generated a point mutant of BRME1 that specifically targets its MEILB2-binding site. The hydrophobic core of the four-helical bundle comprises highly conserved BRME1 amino acids V548, L555, and I562, which occupy the ‘a’ positions within the heptad repeats (Figure 3a). Therefore, we introduced glutamate mutations at these residues (V548E, L555E, and I562E; hereafter referred to as 3E) to disrupt the assembly of the four-helical bundle. We confirmed this disruption biochemically, demonstrating that the 3E mutation effectively prevented the formation of the MEILB2-BRME1 core complex *in vitro* (Figure 3b and Supplementary Figure 3).

**Figure 3.**
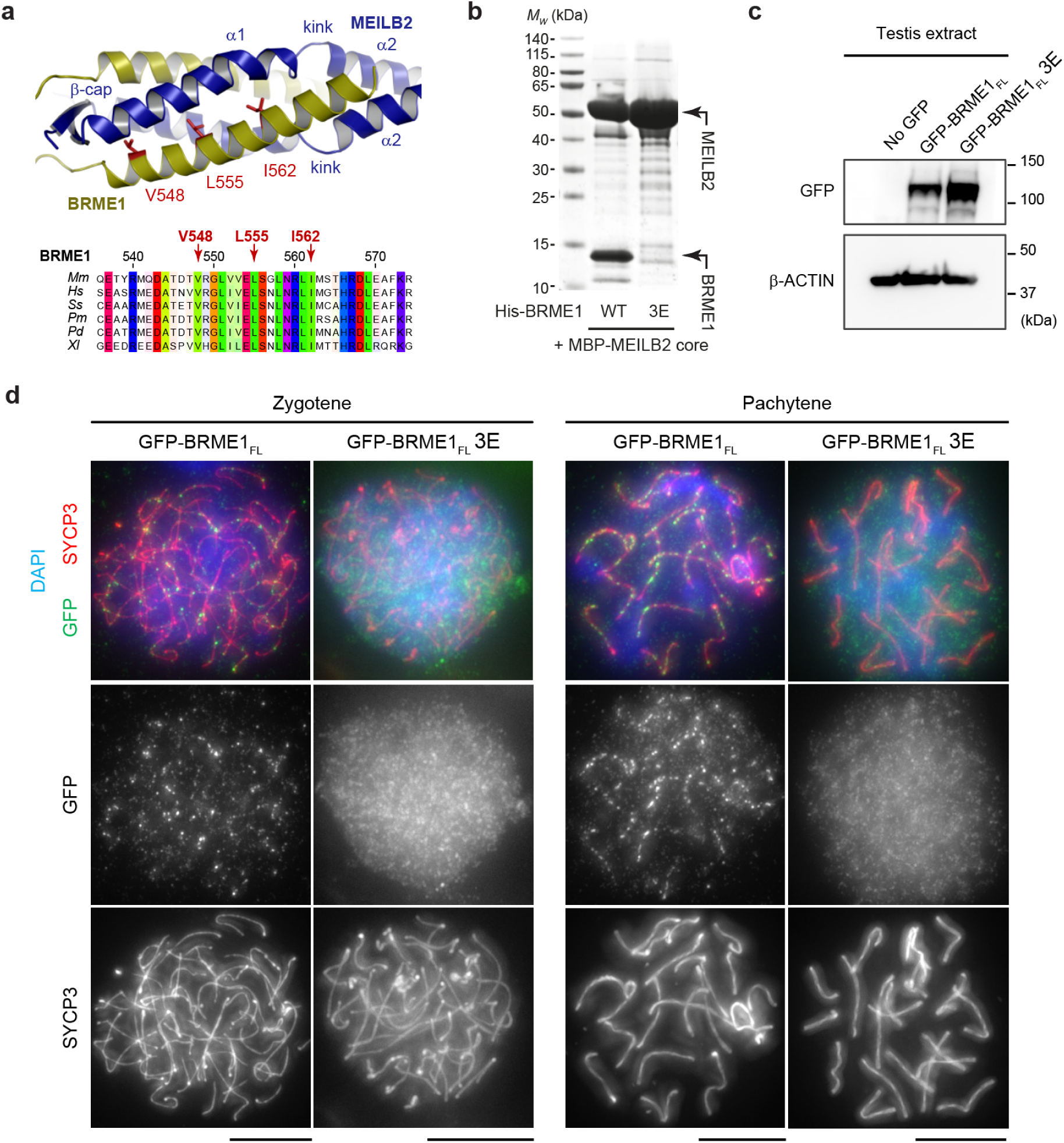
The MEILB2-binding interface is required for BRME1 recruitment to meiotic DSBs. (**a**) The MEILB2-BRME1 interaction involves BRME1 amino-acids V548, L555 and I562, which are buried in the hydrophobic core of the four-helical bundle (top), and are evolutionarily conserved (bottom). Multiple sequence alignment: *Mus musculus* (*Mm*), *Homo sapiens* (*Hs*), *Sus scrofa (Ss)*, *Physeter macrocephalus* (*Pm*), *Phyllostomus discolor* (*Pd*), and *Xenopus laevis* (*Xl*). (**b**) Amylose pull-down of His-BRME1 with MBP-MEILB2 core following recombinant co-expression. MEILB2-binding of BRME1 wild-type (WT) is blocked by the BRME1 V548E L555E I562E (3E) mutation. Controls are shown in Supplementary Figure 3. (**c**,**d**) Expression of GFP-BRME1_FL_ (full-length, wild-type) and GFP-BRME1_FL_ 3E (full-length, 3E mutant) in mouse spermatocytes by *in vivo* electroporation. (**c**) Immunoblots of testis extracts following electroporation with GFP-BRME1_FL_ and GFP-BRME1_FL_ 3E, stained with antibodies against GFP (top) and β-actin (bottom). (**d**) Mouse zygotene (left) and pachytene (right) spermatocytes expressing GFP-BRME1_FL_ and GFP-BRME1_FL_ 3E, stained with anti-SYCP3 antibody (red), anti-GFP antibody (green), and 4,6-diamidino-2-phenylindole (DAPI). Scale bars, 5 μm.

In a previous study, we established that GFP-BRME1 is recruited to meiotic DSBs upon expression in mouse spermatocytes through *in vivo* electroporation (Zhang et al., 2020). Therefore, we employed the same system to investigate the impact of the 3E mutation on the localization of full-length BRME1. While GFP-BRME1_FL_ 3E was expressed at a comparable level to the wild-type protein (Figure 3c), it failed to be recruited to meiotic DSBs in zygotene and pachytene spermatocytes (Figure 3d). These findings confirm that the MEILB2-binding interface of BRME1 observed in the crystal structure is solely responsible for its interaction with MEILB2 and its recruitment to meiotic DSBs *in vivo*.

### Structure of the full MEILB2-BRME1 complex

Our MEILB2-BRME1 core crystal structure contains the N-terminal end of the MEILB2 α2 coiled-coil, up to amino-acid Q77 (Figure 2b). The C-terminal end of the same coiled-coil, from the equivalent of amino-acid Q109, was observed in previous crystal structures of the BRCA2-MEILB2 ARM complex (PDB accessions 7LDG and 7BDX; Pendlebury et al., 2021, Ghouil et al., 2021). Hence, MEILB2 α2 likely forms a continuous coiled-coil that physically separates the BRME1- and BRCA2-binding regions of MEILB2. To visualise this, and build the intervening 31 residues, we generated *Alphafold2* models of the 2:2 complex between the full structured region of MEILB2 (amino-acids 22-338; herein referred to as MEILB2) and BRME1 (Evans et al., 2021, Jumper et al., 2021). During modelling, we enforced the use of our MEILB2-BRME1 core crystal structure and the previous BRCA2-MEILB2 ARM structure (PDB accession 7LDG; Pendlebury et al., 2021) as the sole templates. The resultant model demonstrates that the MEILB2 α2 coiled-coil can seamlessly connect the BRME1-and BRCA2-binding regions of the MEILB2 dimer (Figure 4a and Supplementary Figure 4). Further, it predicts an overall length of approximately 18 nm between the N-terminal β-cap and the C-terminal ARM domains.

**Figure 4.**
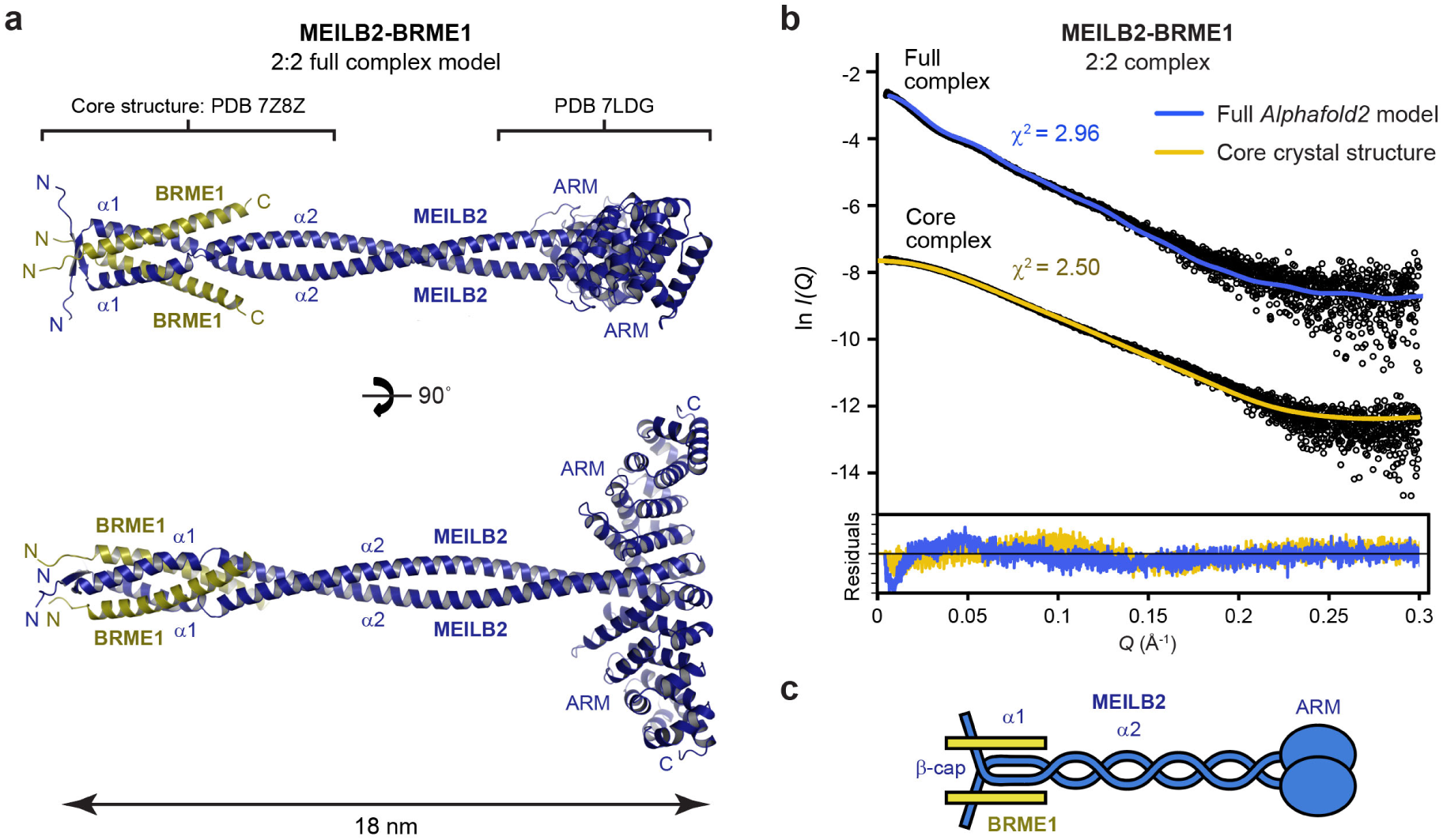
Figure 4 Structure of the full MEILB2-BRME1 2:2 complex. (**a**) *Alphafold2* model of the full MEILB2-BRME1 2:2 complex, generated using the MEILB2-BRME1 2:2 core structure reported herein (PDB accession 7Z8Z), and the MEILB2 ARM domain structure (PDB accession 7LDG; Pendlebury et al., 2021), as the sole templates. The molecule has a length of approximately 18 nm, including a 14-nm separation between the N-terminal four helical bundle and the C-terminal ARM domains. Modelling details, scores and plots are provided in Supplementary Figure 4. (**b**) SEC-SAXS scattering curves of MEILB2-BRME1 2:2 full and core complexes, overlaid with the theoretical scattering curves of the *Alphafold2* model (blue), and the core crystal structure (yellow), with χ^2^ values of 2.96 and 2.50, respectively. Residuals for each fit are shown (inset). Guinier analyses and real-space *P(r)* distributions are shown in Supplementary Figure 5. (**c**) Schematic of the MEILB2-BRME1 2:2 structure.

We confirmed the validity of the MEILB2-BRME1 2:2 model by size-exclusion chromatography small-angle X-ray scattering (SEC-SAXS) analysis of the MEILB2-BRME1 complex in solution. The SAXS real-space *(r)* pair-distance distribution indicated a rod-like molecule of 20-nm length, including a 14-nm separation between domains, consistent with the length and location of four-helical bundle and ARM domains within the model (Figure 4b and Supplementary Figure 5). Further, the SAXS scattering curve was closely fitted by the MEILB2-BRME1 model, similar to the fit between the SAXS data and crystal structure of the core complex, at χ^2^ values of 2.96 and 2.50, respectively (Figure 4b). Hence, we conclude that the MEILB2-BRME1 has a linear structure in which the N-terminal BRME1-binding core is connected to C-terminal ARM domains by a MEILB2 α2 coiled-coil (Figure 4c).

**Figure 5.**
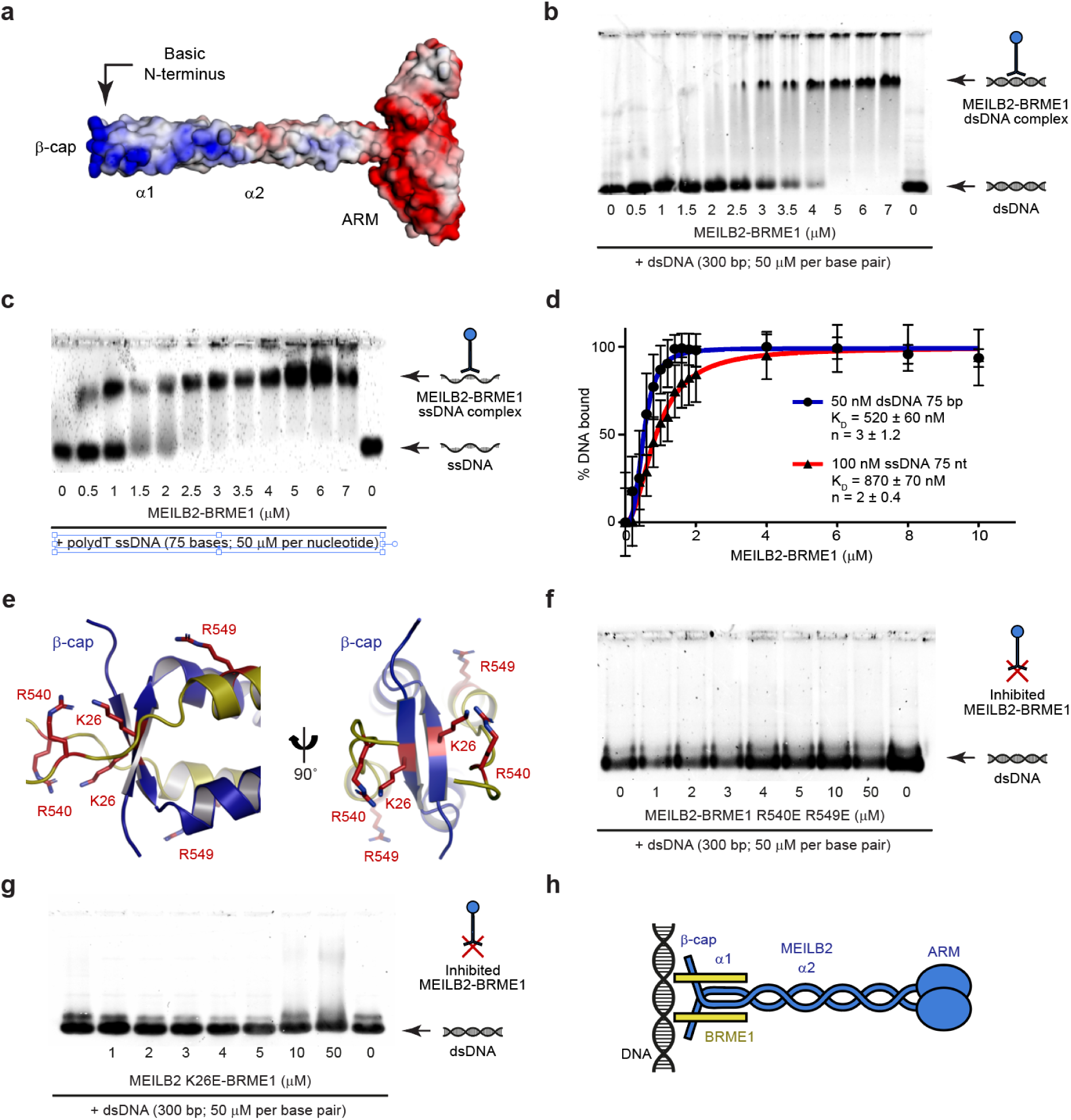
DNA-binding is mediated by the β-cap of MEILB2-BRME1. (**a**) Surface electrostatic potential of the MEILB2-BRME1 modelled structure (red, electronegative; blue, electropositive). (**b,c**) Electrophoretic mobility shift assays (EMSAs) analysing the ability of MEILB2-BRME1 (at molecular concentrations indicated) to interact with (**b**) dsDNA (300 base pair) and (**c**) polydT ssDNA (75 bases), at concentrations of 50 μM per nucleotide/base pair. (**d**) Quantification of DNA-binding by MEILB2-BRME1 through densitometry of EMSAs performed using 50 nM (per molecule) FAM-dsDNA (75 base pairs; blue) and 100 nM (per molecule) FAM-ssDNA (75 nucleotides; red). Plots, *K_D_* and n values were determined by fitting data to the Hill equation and are quoted within a 95% confidence interval; error bars indicate standard error, *n* = 3 EMSAs. (**e**) The N-terminal β-cap of the MEILB2-BRME1 2:2 core structure includes basic amino-acids MEILB2 K26 and BRME1 R540 and R549. (**f**,**g**) EMSAs analysing the ability of MEILB2-BRME1 (at molecular concentrations indicated) harbouring (**f**) BRME1 R540E R549E and (**g**) MEILB2 K26E mutations to interact with dsDNA (300 base pair) at concentrations of 50 μM per base/base pair. (**c,d,f,g**) Gel images are representative of at least three replicates performed using different protein samples. (**h**) Schematic of MEILB2-BRME1 interacting via its β-cap with DNA.

### MEILB2-BRME1 binds to DNA via its β-cap

The surface electrostatics of the MEILB2-BRME1 2:2 model indicated that the N-terminal MEILB2-BRME1 core contains discrete patches of basic charge, whereas the C-terminal ARM domains are predominantly acidic (Figure 5a). Hence, we wondered whether the N-terminal end of the molecule may bind to DNA. Accordingly, electrophoretic mobility assays (EMSAs) showed that MEILB2-BRME1 forms discrete protein-DNA complexes with dsDNA and poly-dT ssDNA (Figure 5b,c). DNA-binding showed no overt sequence specificity, and saturation occurred at ratios of one MEILB2-BRME1 2:2 complex to 10 base pairs (dsDNA) or 20 nucleotides (ssDNA) (Figure 5b,c). Quantification of EMSAs conducted with substrate concentrations of 50 nM and 100 nM, and fitting to the Hill equation, revealed positive cooperativity with binding affinities of 520 ± 60 nM and 870 ± 70 nM for dsDNA and ssDNA, respectively (Figure 5d). These findings are consistent with MEILB2-BRME1 binding to the DNA backbone, with a binding footprint spanning ten base pairs.

Notably, the basic charge of MEILB2-BRME1 is particularly concentrated at its N-terminal β-cap (Figure 5a). Given that the β-cap stabilises the end of the coiled-coil, we hypothesised that it may serve as a rigid platform for DNA binding at the tip of the MEILB2-BRME1 complex. To test this, we introduced glutamate mutations of MEILB2 amino-acid K26, and BRME1 amino-acids R540 and R549, which are the main contributors of basic charge at the β-cap. Consequently, DNA binding of MEILB2-BRME1 was abolished by both the MEILB2 K26E and BRME1 K540E R549E mutations (Figure 5f,g). Thus, DNA-binding is mediated by the rigid β-cap at the N-terminal tip of the MEILB2-BRME1 structure (Figure 5h). Further, DNA-binding involves amino-acids from both MEILB2 and BRME1, so is a consequence of complex formation rather than being an intrinsic property of either component.

### BRCA2 dimerises MEILB2-BRME1 to form a V-shaped assembly

What is the structure of the complex formed by MEILB2-BRME1 upon interaction with BRCA2? It was previously shown that BRCA2-binding dimerises ARM domains from opposing MEILB2 dimers to form a stable 2:4 complex (Pendlebury et al., 2021, Ghouil et al., 2021). To test whether this also occurs in the presence of BRME1, we used size-exclusion chromatography multi-angle light scattering (SEC-MALS) to determine the oligomeric state of MEILB2-BRME1 upon binding to BRCA2’s MBD (amino-acids 2232-2283; herein referred to as BRCA2). This revealed that MEILB2-BRME1 and BRCA2-MEILB2-BRME1 are stable 2:2 and 2:4:4 complexes, respectively (5Figure 5a). Hence, BRCA2-binding induces dimerization of MEILB2-BRME1 to form a stable 2:4:4 ternary complex.

We next modelled the BRCA2-MEILB2-BRME1 2:4:4 structure by docking two MEILB2-BRME1 models onto the constituent MEILB2 ARM dimers of the previous BRCA2-MEILB2 ARM 2:4 crystal structure (PDB accession 7LDG; Pendlebury et al., 2021) (Supplementary Figure 6). The resultant model reveals a V-shaped assembly in which MEILB2-BRME1 2:2 complexes are held at approximately 90° to one another, with their opposing ARM domains stapled together by BRCA2 MBDs (Figure 6b). The unusual architecture of this assembly imposes a separation of approximately 25 nm between the β-caps at the tips of its limbs (Figure 6b). To validate the model, the solution structure of the BRCA2-MEILB2-BRME1 2:4:4 complex was analysed using SEC-SAXS. The SAXS real-space P(r) pair-distance distribution indicated an elongated molecule with a maximum dimension of 27 nm, consistent with the predicted geometry of the model (Figure 6c and Supplementary Figure 7). Further, the SAXS scattering curve was closely fitted by the 2:4:4 model (*χ*^2^ = 1.36; Figure 6c). Hence, BRCA2-binding induces dimerization of MEILB2-BRME1 2:2 complexes into a V-shaped 2:4:4 assembly, where the β-caps at the N-terminal tips of its two limbs are separated by approximately 25 nm (Figure 6d).

**Figure 6.**
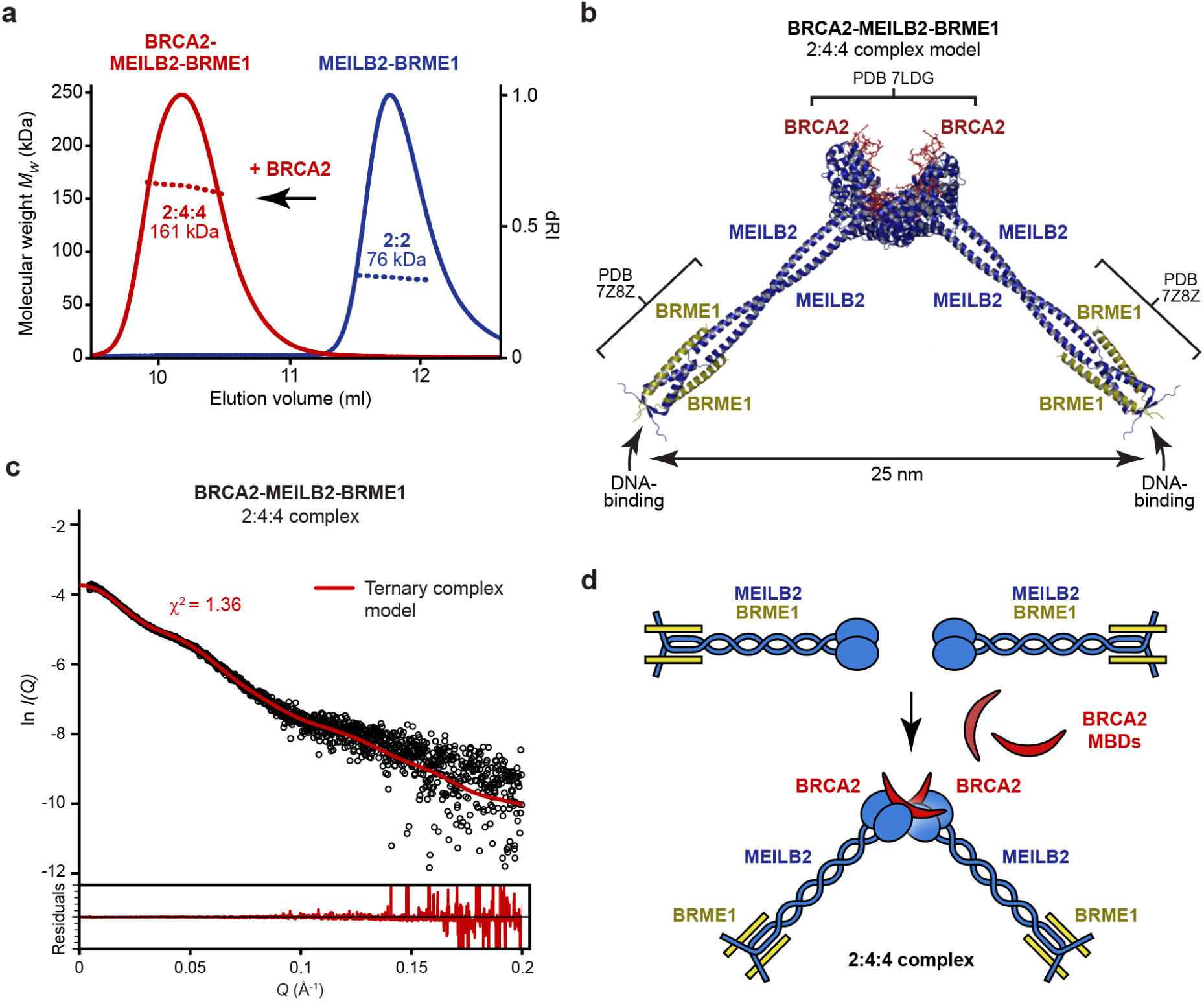
Structure of the BRCA2-MEILB2-BRME1 2:4:4 ternary complex. (**a**) SEC-MALS analysis of MEILB2-BRME1 (blue) and the BRCA2-MEILB2-BRME1 ternary complex (red), showing differential refractive index (dRI; solid lines) profiles with fitted molecular weights (*Mw*; dashed lines) across elution peaks. MEILB2-BRME1 forms a 76 kDa 2:2 complex (theoretical *Mw* – 82 kDa), and BRCA2-MEILB2-BRME1 forms a 161 kDa 2:4:4 complex (theoretical *Mw* – 176 kDa). (**b**) Model of the BRCA2-MEILB2-BRME1 2:4:4 complex generated by docking two MEILB2-BRME1 2:2 complex models (shown in 5Figure 4a) onto the BRCA2-MEILB2 ARM domain 2:4 complex crystal structure (PDB accession 7LDG; Pendlebury et al., 2021). The model predicts that the DNA-binding β-caps of the two constituent MEILB2-BRME1 core complexes are separated by 25 nm. Modelling details are shown in Supplementary Figure 6. (**c**) SEC-SAXS data in which the BRCA2-MEILB2-BRME1 scattering cure is overlaid with the theoretical scattering curve of the modelled structure (red; shown in panel b), with a χ^2^ value of 1.36. Residuals are shown (inset). Guinier analysis and real-space *P(r)* distribution are shown in Supplementary Figure 7. (**d**) Schematic of BRCA2-MEILB2-BRME1 complex formation, in which BRCA2-binding induces dimerization of two MEILB2-BRME1 2:2 complexes into a V-shaped 2:4:4 complex.

### BRCA2-MEILB2-BRME1 acts as a DNA clamp

The physical separation between the two DNA-binding β-caps of the BRCA2-MEILB2-BRME1 2:4:4 ternary complex suggested that may be able to bridge between DNA molecules. Accordingly, EMSAs demonstrated that BRCA2-MEILB2-BRME1 readily bound to a 75 base-pair dsDNA substate, forming protein-DNA networks that were too large to enter the gel (Figure 7a). To temper DNA-binding, and hence aid visualisation of protein-DNA species, we also analysed a BRCA2-MEILB2-BRME1 complex in which BRME1 was truncated to remove basic C-terminal amino-acids (deletion of 575-RKTK-579; herein referred to as BRME1 ΔRKTK). This slightly diminished DNA-binding and reduced protein-DNA networks to sizes that were resolved by EMSA (Figure 7b).

**Figure 7.**
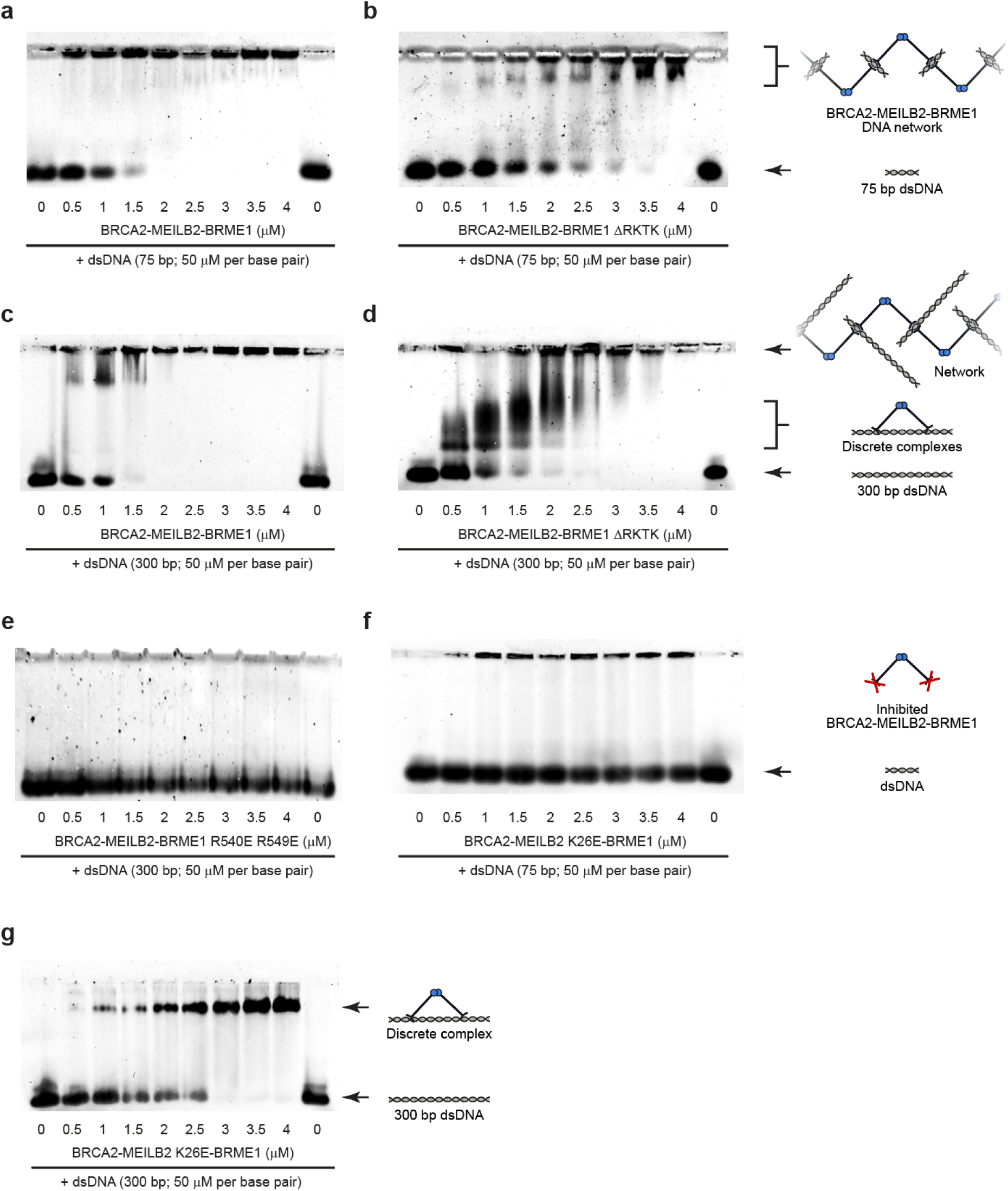
BRCA2-MEILB2-BRME1 bridges between DNA molecules. (**a-d**) EMSAs analysing the ability of (**a,c**) BRCA2-MEILB2-BRME1 and (**b,d**) BRCA2-MEILB2-BRME1 ΔRKTK (truncation of BRME1 to remove the last four amino-acids), at molecular concentrations indicated, to interact with (**a,b**) 75 base pair dsDNA and (**c,d**) 300 base pair dsDNA, at concentrations of 50 μM per base pair. (**e-g**) EMSAs analysing the ability of BRCA2-MEILB2-BRME1 harbouring (**e**) BRME1 R540E R549E and (**f,g**) MEILB2 K26E mutations, at molecular concentrations indicated, to interact with (**e,g**) 300 base pair dsDNA and (**f**) 75 base pair dsDNA, at concentrations of 50 μM per base pair.

We reasoned that if the DNA substrate were of sufficient length, it should be possible for the two β-caps of the ternary complex to bind cooperatively to the same DNA molecule. Hence, we analysed a 300 base-pair substrate, which has a length of 90 nm that far exceeds the 25 nm gap between β-caps. EMSAs showed that BRCA2-BRME1-MEILB2 initially formed discrete species with the 300 base-pair substrate, and then generated protein-DNA networks when present at stoichiometric excess (Figure 7c). This effect was particularly pronounced for the tempered BRCA2-BRME1-MEILB2 ΔRKTK complex, which formed dominant discrete species prior to assembly at higher ratios into networks (Figure 7d). These discrete intermediate species were not present for the 75 base-pair substrate (Figure 7a,b), which has a length of 22 nm, so is insufficient to span between the two DNA-binding β-caps. These data strongly support a model in which the two β-caps of the ternary complex act as spatially separated DNA-binding sites that can bridge between discrete DNA molecules.

The BRME1 R540 R549E mutation abrogated DNA-binding of the BRCA2-MEILB2-BRME1 complex (Figure 7e), confirming that DNA-binding of the ternary complex is mediated by the β-caps. The MEILB2 K26E mutation also blocked binding of the BRCA2-MEILB2-BRME1 complex to the shorter 75 base-pair substrate (Figure 7f). However, it retained DNA-binding to the longer 300 base-pair substrate, albeit forming a single discrete species rather than protein-DNA networks (Figure 7g). Hence, whilst the MEILB2 K26E mutation was sufficient to block DNA-binding by a single β-cap, it likely retained sufficient residual affinity for the two β-caps of the ternary complex to interact cooperatively with a sufficiently long single DNA molecule. Thus, the MEILB2 K26E mutant provides additional evidence in favour of our model.

In summary, BRCA2-binding induces dimerization of MEILB2-BRME1 into a V-shaped 2:4:4 assembly, in which DNA-binding β-caps at the tips of its two limbs can independently interact with DNA molecules. Hence, BRCA2 induces MEILB2-BRME1 to act as a DNA clamp that holds together discrete DNA molecules, with a separation of approximately 25 nm (Figure 8). Thus, we propose that BRCA2 induces the clamping together of DNA molecules by MEILB2-BRME1 to facilitate inter-homologue recombination as part of the specialised mechanics of meiotic recombination.

**Figure 8.**
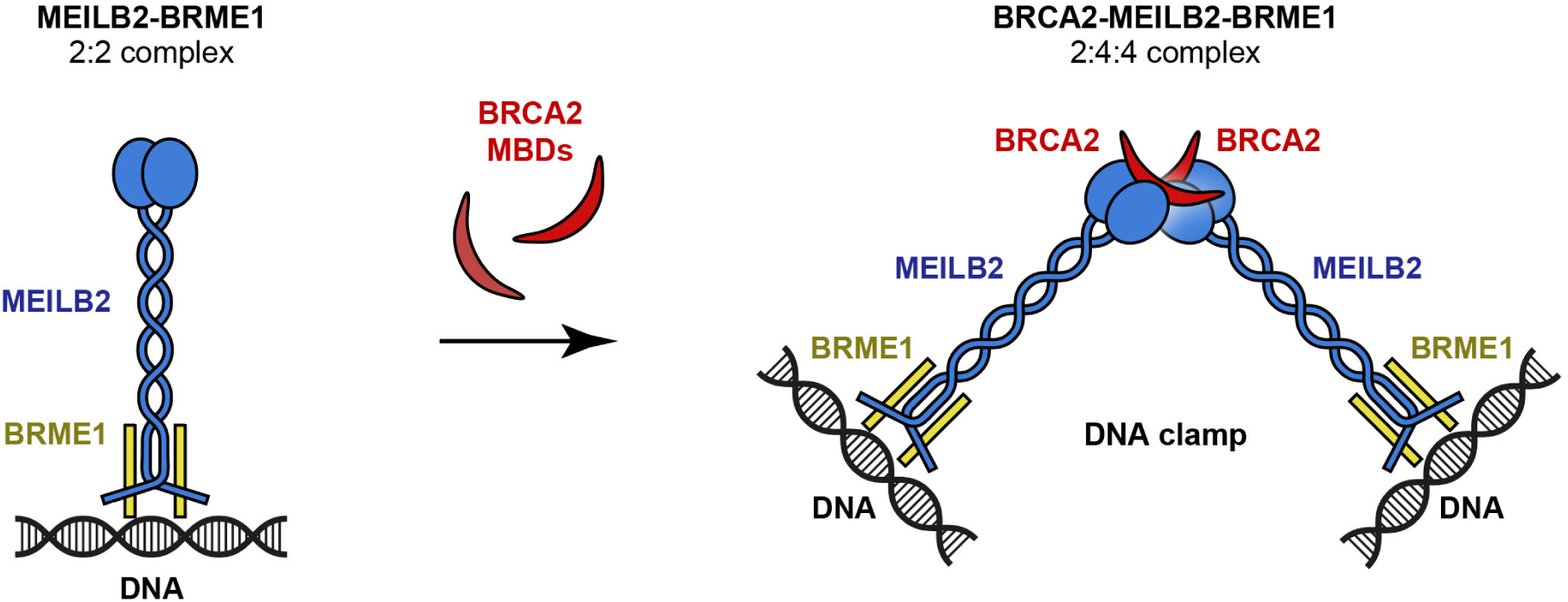
Model of BRCA2-induced dimerization of MEILB2-BRME1 into a V-shaped DNA clamp. MEILB2-BRME1 2:2 complexes bind to DNA via their N-terminal β-caps. BRCA2-binding induces dimerization of MEILB2-BRME1 2:2 complexes into a V-shaped 2:4:4 assembly, in which β-caps are physically separated by 25 nm and can clamp together DNA molecules. This could bridge between resected ends of the meiotic DSB, or between homologous chromosomes. The BRCA2-induced MEILB2-BRME1 DNA clamp may act in concert with the proximal DNA-binding domain and RAD51-/DMC1-binding regions of BRCA2, in a coordinated assembly to fulfil the specialised requirements of meiotic recombination.

## Discussion

The molecular programme of meiosis utilises the machinery of DNA double-strand break repair by homologous recombination to align and generate crossovers between homologous chromosomes that are critical for their correct segregation and the formation of haploid germ cells. The particular adaptations of meiotic recombination require several additional meiosis-specific components. These include the MEILB2-BRME1 complex, which binds directly to BRCA2 and is essential for the correct localisation of RAD51 and DMC1 recombinases to meiotic DSBs. Here, we combine crystallographic and cell biology data to report the structure of the core MEILB2-BRME1 2:2 complex, which we combine with existing structural data to generate an experimentally validated model of BRCA2-MEILB2-BRME1 2:4:4 ternary complex. This V-shaped structure contains DNA-binding sites at the tips of its two limbs, which are separated by 25 nm, suggesting that it may function to clamp together DNA molecules. As BRCA2-binding is necessary for dimerization of MEILB2-BRME1 into the V-shaped structure, we conclude that its proposed role as a DNA clamp occurs within the context of its complex with BRCA2 (Figure 8).

What is the function of the BRCA2-induced MEILB2-BRME1 DNA clamp in meiosis? We envisage two possibilities. Firstly, it may bridge between DSB ends, in a manner that favours their repair by inter-homologue recombination. This is reminiscent of how MRN and CtIP are proposed to tether DNA ends and affect the choice of repair pathway at DSB sites (Oh and Symington, 2018, Davies et al., 2015, Wilkinson et al., 2019). Alternatively, it may bridge between homologous chromosomes to provide stabilising interactions between invading RAD51/DMC1-ssDNA nucleoprotein filaments and the template during inter-homologue recombination. This is an attractive model as it fulfils a unique requirement of meiosis, which is provided in mitotic recombination by cohesion between sister chromatids (Piazza et al., 2021). Notably, the 25 nm distance between DNA-binding sites of the dimerised MEILB2-BRME1 complex is comparable to the approximately 30 nm diameter of cohesin rings (Nasmyth, 2005). Hence, BRCA2-induced MEILB2-BRME1 DNA clamps may facilitate inter-homologue recombination by bridging between homologous chromosomes in the same manner as cohesin facilitates inter-sister recombination by tethering together sister chromatids.

A particular requirement of meiotic recombination is the removal of MEIOB2-SPATA22, in addition to RPA, from 5’-ssDNA overhangs for loading of RAD51 and DMC1 recombinases (Souquet et al., 2013, Ribeiro et al., 2021, Luo et al., 2013). As BRCA2-DSS1 can offload RPA (Bell et al., 2023), it was proposed that MEILB2-BRME1 may displace MEIOB2-SPATA22 during meiotic recombination (Zhang et al., 2022). This is supported by the co-immunoprecipitation of MEIOB-SPATA22 with MEILB2, and the accumulation of MEIOB-SPATA22 at DSBs in MEILB2 and BRME1 knockouts (Zhang et al., 2019, Zhang et al., 2020, Takemoto et al., 2020). Our finding that MEILB2-BRME1 binds directly to DNA, with no overt specificity for single- or double-stranded substrates, is consistent with this proposed role. Indeed, as BRCA2’s MBD is proximal to its globular DNA-binding domain, we speculate that BRCA2-DSS1 and MEILB2-BRME1 may act together, in a coordinated manner, to offload RPA and MEIOB2-SPATA22 during meiotic recombination. This may occur alongside its proposed role in bridging between DNA ends or homologous chromosomes.

Thus far, we have addressed how BRCA2-binding affects MEILB2-BRME1 structure, but we must also consider the converse issue of how MEILB2-BRME1 affects the structure of BRCA2. In isolation, BRCA2 can form monomers, dimers and larger oligomers, in a manner that involves its N-terminal and C-terminal regions, and is modulated by ssDNA, DSS1 and RAD51 (Le et al., 2020, Paul et al., 2021). Further, the BRCA2 dimer undergoes conformational change upon binding to RAD51 (Shahid et al., 2014). In meiosis, binding to MEILB2-BRME1 would clearly dimerise BRCA2 through its MBD region. However, given the lack of high-resolution structures of BRCA2 oligomers, it is not possible to say whether this would reinforce existing dimers or form alternative oligomers. In addition, binding to MEILB2-BRME1 may alter the conformation of BRCA2, such as to affect DMC1-binding and DNA-binding of the sites immediately downstream of the MBD. Any effects on the oligomeric state and conformation of BRCA2 likely favour inter-homologue rather than inter-sister recombination (Zhang et al., 2022). Hence, these interactions are likely to be deleterious if they occur outside of meiosis. Accordingly, MEILB2 is expressed in cancer cell lines and human tumour samples (Brandsma et al., 2019), and ectopic expression of MEILB2 inhibits homologous recombination in somatic cells (Sato et al., 2020, Zhang et al., 2020). Whilst BRME1 is also upregulated in cancers and suppresses recombination upon ectopic expression (Zhang et al., 2020), it is not yet possible speculate how it affects recombination as the structure and function of all but the MEILB2-binding site are unknown.

Overall, we propose that BRCA2-binding of MEILB2-BRME1 may clamp together resected ends or homologous chromosomes, may facilitate the removal of MEIOB-SPATA22 from 5’-ssDNA overhangs, and may alter the oligomeric state and/or conformation of BRCA2. These are not mutually exclusive, so may represent multifactorial roles of MEIB2-BRME1 in meiotic recombination. Further, MEILB2-BRME1 likely acts in coordination with the DMC1-binding site and globular DNA-binding domain that are immediately downstream of the MBD. Hence, the ternary complex formed by MEILB2-BRME1, BRCA2-DSS1, RAD51 and DMC1 may operate as a single functional unit that fulfils the above roles to facilitate inter-homologue recombination. Elucidating the structure of this meiotic ‘recombinosome’ will ultimately reveal the molecular mechanisms that underpin meiotic recombination. Further, the unusual architecture imposed by MEILB2-BRME1 may provide unique structural insights that will uncover the molecular basis of the wider role of BRCA2 in recombination-mediated DNA repair.

## Materials and methods

### Recombinant protein expression and purification

Constructs of mouse MEILB2 (amino acid residues: 22-81, 22-338) and mouse BRCA2 (amino acid residues: 2232-2283) were cloned into pMAT11 (Peranen et al., 1996) and pRSF-Duet1 (Merck Millipore) vectors for expression with an N-terminal TEV-cleavable His_6_-MBP tag and an N-terminal TEV-cleavable MBP tag, respectively. Constructs of mouse BRME1 (amino acid residues: 540-578 and 540-574) were cloned into pMAT11 (Peranen et al., 1996) and pRSF-Duet1 (Merck Millipore) vectors for expression with an N-terminal TEV-cleavable His_6_ tag. Protein constructs were expressed in BL21 (DE3) cells (Novagen), in 2xYT media, and induced with 0.5 mM IPTG for 16 h at 25°C. Bacterial pellets were harvested, resuspended in 20 mM Tris, pH 8.0, 500 mM KCl, and lysed using a TS Cell Disruptor (Constant Systems) at 172 MPa. Cellular debris was later removed by centrifugation at 40,000 g. Fusion proteins were purified through consecutive Ni-NTA (Qiagen), amylose (NEB), and HiTrap Q HP (Cytiva) ion exchange chromatography. The N-terminal tags were cleaved using TEV protease, and the cleaved samples were further purified through HiTrap Q HP (Cytiva) ion exchange chromatography and size exclusion chromatography (HiLoad 16/600 Superdex 200, Cytiva) in 20 mM HEPES, pH 7.5, 150 mM KCl, 2 mM DTT for MEILB2-BRME1 constructs and in 20 mM HEPES, pH 7.5, 300 mM KCl, 2 mM DTT for BRCA2-MEILB2-BRME1 constructs. Purified protein samples were spin-concentrated using Amicon Ultra centrifugal filter device (10,000 NMWL), flash-frozen in liquid nitrogen, and stored at −80°C. Purified proteins were analysed using SDS-PAGE and visualized with Coomassie staining. Protein concentrations were determined using Cary 60 UV spectrophotometer (Agilent) with extinction coefficients and molecular weights calculated by ProtParam (http://web.expasy.org/protparam/).

### Crystal structure of MEILB2-BRME1 core (PDB accession 7Z8Z)

MEILB2-BRME1 (22-81; 540-574) protein crystals grew upon incubation of protein at ∼10 mg/ml on ice, in 20mM HEPES pH 7.5, 150mM KCl. Crystals were cryo-protected by addition of 30% PEG 400, and were cryo-cooled in liquid nitrogen. X-ray diffraction data were collected at 0.9795 Å, 100 K, as four separate datasets, each of 3600 consecutive 0.10° frames of 0.020 s exposure on a Dectris Eiger2 XE 16M detector at beamline I04 of the Diamond Light Source synchrotron facility (Oxfordshire, UK). Data were indexed, integrated in XDS (Kabsch, 2010), scaled and merged in Aimless (Evans, 2011), using AutoPROC (Vonrhein et al., 2011). Crystals belong to orthorhombic spacegroup P2_1_2_1_2_1_ (cell dimensions a = 27.05 Å, b = 43.99 Å, c = 178.71 Å, α = 90°, β = 90°, γ = 90°), with one MEILB2-BRME1 2:2 complex in the asymmetric unit. Structure solution was achieved through fragment-based molecular replacement using ARCIMBOLDO_LITE (Rodriguez et al., 2009), in which eight helices of 14 amino-acids were placed by PHASER (McCoy et al., 2007) and extended by tracing in SHELXE utilising its coiled-coil mode (Caballero et al., 2018). A correct solution was identified by a SHELXE correlation coefficient of 46.6%. Model building was performed through iterative re-building by PHENIX Autobuild (Adams et al., 2010) and manual building in Coot (Emsley et al., 2010). The structure was refined using PHENIX refine (Adams et al., 2010), using anisotropic (protein) and isotropic (water) atomic displacement parameters. The structure was refined against data to an isotropic resolution limit of 1.50 Å, to R and R_free_ values of 0.1965 and 0.2347, respectively, with 100% of residues within the favoured regions of the Ramachandran plot (0 outliers), clashscore of 1.85 and overall MolProbity score of 0.95 (Chen et al., 2010).

### Size-exclusion chromatography multi-angle light scattering (SEC-MALS)

The absolute molar masses of protein samples were determined by multiangle light scattering coupled with size exclusion chromatography (SEC-MALS). Protein samples at >5 mg/ml were loaded onto a Superdex 200 Increase 10/300 GL size exclusion chromatography column (Cytiva) in 20 mM HEPES, pH 7.5, 150 mM KCl, 2 mM DTT for MEILB2-BRME1 constructs and in 20 mM HEPES, pH 7.5, 500 mM KCl, 2 mM DTT for BRCA2-MEILB2-BRME1 constructs, at 0.5 ml/min, in line with a DAWN HELEOS II MALS detector (Wyatt Technology) and an Optilab T-rEX differential refractometer (Wyatt Technology). Differential refractive index and light scattering data were collected and analyzed using

ASTRA 6 software (Wyatt Technology). Molecular weights and estimated errors were calculated across eluted peaks by extrapolation from Zimm plots using a dn/dc value of 0.1850 ml/g. Bovine serum albumin (Thermo Fisher Scientific) was used as the calibration standard.

### Size-exclusion chromatography small-angle X-ray scattering (SEC-SAXS)

SEC-SAXS experiments were performed at beamline B21 of the Diamond Light Source synchrotron facility (Oxfordshire, UK). Protein samples at concentrations >8 mg/ml were loaded onto a Superdex™ 200 Increase 10/300 GL size exclusion chromatography column (GE Healthcare) in 20 mM HEPES pH 7.5, 150 mM KCl, 2 mM DTT for MEILB2-BRME1 constructs and in 20 mM HEPES pH 7.5, 500 mM KCl, 2 mM DTT for BRCA2-MEILB2-BRME1 constructs at 0.5 ml/min using an Agilent 1200 HPLC system. The column outlet was fed into the experimental cell, and SAXS data were recorded at 12.4 keV, detector distance 4.014 m, in 3.0 s frames. ScÅtter 3.0 was used to subtract, average the frames and carry out the Guinier analysis for the radius of gyration (*Rg*), and *P(r)* distributions were fitted using *PRIMUS* (P.V.Konarev, 2003). Crystal structures and models were fitted to experimental data using *CRYSOL* (Svergun D.I., 1995).

### Electrophoretic mobility shift assays (EMSAs)

Protein complexes were incubated with 50 μM (per base pair) 75 bp and 300bp linear random sequence dsDNA, and 75 nt FAM-poly(dT) ssDNA, at concentrations indicated, in 20 mM HEPES pH 7.5, 325 mM KCl for 60 mins at room temperature. Glycerol was added at a final concentration of 3%, and samples were analysed by electrophoresis on a 0.4% (w/v) agarose gel in 0.5x TBE pH 8.0 at 20-40 V for ∼4 h at 4 °C. DNA was detected by SYBR™ Gold (ThermoFisher).

### *K_D_* determination by EMSA

Quantification of DNA-binding was performed though EMSA (as described above) using 50 nM FAM-labelled 75 bp random sequence dsDNA and 100 nM 75 nt poly(dT), at protein concentrations indicated. DNA was detected by FAM and SYBR™ Gold (ThermoFisher) staining using a ChemiDoc MP Imaging System (Bio-Rad). Gels were analysed using Image Lab software (Bio-Rad). The DNA-bound proportion was plotted against molecular protein concentration and fitted to the Hill–Langmuir equation (below), with apparent *K_D_* determined, using Prism8 (GraphPad). Protein concentrations used for *K_D_* estimation are quoted for the oligomeric species.

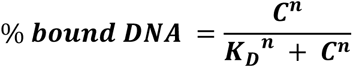

### Structural modelling of the full MEILB2-BRME1 2:2 complex

Models were generated using a local installation of *Alphafold2* v2.3.2 (Jumper et al., 2021) that was modified to control the use of PDB structures and newly solves crystal structures as templates. Models of the full MEILB2-BRME1 2:2 complex were generated through the multimer pipeline (Evans et al., 2021), using the MEILB2-BRME1 2:2 core crystal structure 7Z8Z reported herein, and PDB structure 7LDG (Pendlebury et al., 2021), as the sole templates. Modelling data were analysed using modules from the ColabFold notebook (Mirdita et al., 2022).

### Structural modelling of the BRCA2-MEILB2-BRME1 2:4:4 complex

A model of the BRCA2-MEILB2-BRME1 2:4:4 complex was generated by docking two MEILB2-BRME1 2:2 complex models onto opposing MEILB2 ARM domain dimers of the BRCA2-MEILB2 ARM 2:4 crystal structure (PDB accession 7LDG; Pendlebury et al., 2021). The docked MEILB2-BRME1 2:2 complexes and BRCA2 chains from the structure were then combined. Models were generated using *PyMOL* Molecular Graphics System, Version 2.4 Schrödinger, LLC.

### Protein sequence and structure analysis

MEILB2 and BRME1 sequences were aligned and visualised using MUSCLE (Madeira et al., 2019) and Jalview (Waterhouse et al., 2009). Molecular structure images were generated using the PyMOL Molecular Graphics System, Version 2.4 Schrödinger, LLC.

### Mice

Wild type mice were congenic with the C57BL/6J background. The mice are housed in IVC cages with a 12 hour dark and light cycle. The temperature is 20-22 °C and the relative humidity is between 45 and 60 %. The mice have bedding material in the form of wood shavings and wood litter as well as a house of paper and nesting pads as enrichment. Cage changing is done at least once a week. All animal experiments were approved by the Regional Ethics Committee of Gothenburg, governed by the Swedish Board of Agriculture (#1316/18).

### Antibodies

The following antibodies were used: rabbit antibodies against GFP (Invitrogen; A11122, 2339829); mouse antibodies against β-Actin (Sigma; A2228-200UL, 067M4856V); and chicken antibody against SYCP3 (Shibuya lab).

### Exogenous protein expression in the testis

Plasmid DNA was electroporated into live mouse testes as previously described (Shibuya et al., 2014). Briefly, male mice at postnatal day 16–20 were anesthetized with pentobarbital, and the testes were pulled from the abdominal cavity. Plasmid DNA (10 μl of 5 μg/μl solution) was injected into each testis using glass capillaries under a stereomicroscope. Testes were held between a pair of tweezers-type electrodes (CUY21; BEX), and electric pulses were applied four times and again four times in the reverse direction at 35 V for 50 ms for each pulse. The testes were then returned to the abdominal cavity, and the abdominal wall and skin were closed with sutures. The testes were removed 24 h after the electroporation, and immunostaining was performed.

### Immunostaining of spermatocytes

Testis cell suspensions were prepared by mincing the tissue with flathead forceps in PBS, washing several times in PBS, and resuspending in a hypotonic buffer (30 mM Tris (pH 7.5), 17 mM trisodium citrate, 5 mM EDTA, 2.5 mM DTT, 0.5 mM PMSF, and 50 mM sucrose). After 30 min, the sample was centrifuged and the supernatant was aspirated. The pellet was resuspended in 100 mM sucrose. After 10 min, an equal volume of fixation buffer (1% paraformaldehyde and 0.1% Triton X-100) was added. Cells were applied to a glass slide, allowed to fix for 2 h at room temperature, and air-dried. For immunostaining, the slides were incubated with primary antibodies in PBS containing 5% BSA for 2 h and then with the following secondary antibodies for 1h at room temperature: Donkey Anti-Rabbit Alexa 488 (1:1000; Invitrogen; A21206, 2376850) and Donkey Anti-Chicken Alexa 594 (1:1000; Invitrogen; A78951, 2551396). The slides were washed with PBS and mounted with VECTASHIELD medium with DAPI (Vector Laboratories).

### Microscopy

Images were obtained on a microscope (Olympus IL-X71 Delta Vision; Applied Precision) equipped with 100× NA 1.40 objective, a camera (CoolSNAP HQ; Photometrics), and softWoRx 5.5.5 acquisition software (Delta Vision). Images were processed with Photoshop (Adobe).

## Data availability

Crystallographic structure factors and atomic coordinates have been deposited in the Protein Data Bank (PDB) under accession number 7Z8Z, and raw diffraction data have been uploaded to https://proteindiffraction.org/.

## Acknowledgements

We thank Diamond Light Source and the staff of beamlines I04 and B21 (proposal mx25233). This work was supported by a Wellcome Senior Research Fellowship to O.R.D. (Grant Number 219413/Z/19/Z), a core grant to the Wellcome Centre for Cell Biology (203149), and the European Research Council (StG-801659).

## Author contributions

M.G. performed all the *in vitro* experimental work. J.Z. and K.Z. performed the *in vivo* experimental work under the supervision of H.S.. O.R.D. designed experiments, analysed data and wrote the manuscript.

---

**Supplementary Figure 1.**
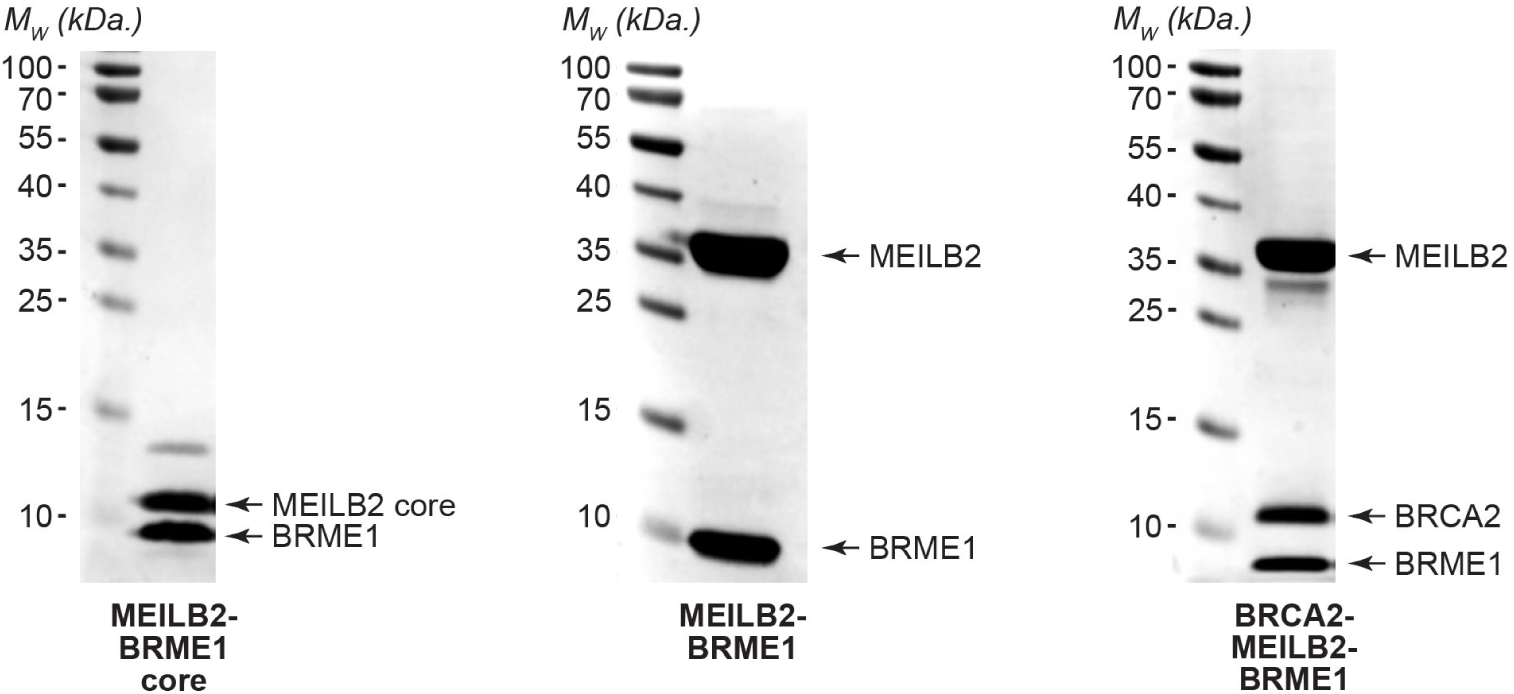
SDS-PAGE of protein samples used in this study.

**Supplementary Figure 2.**
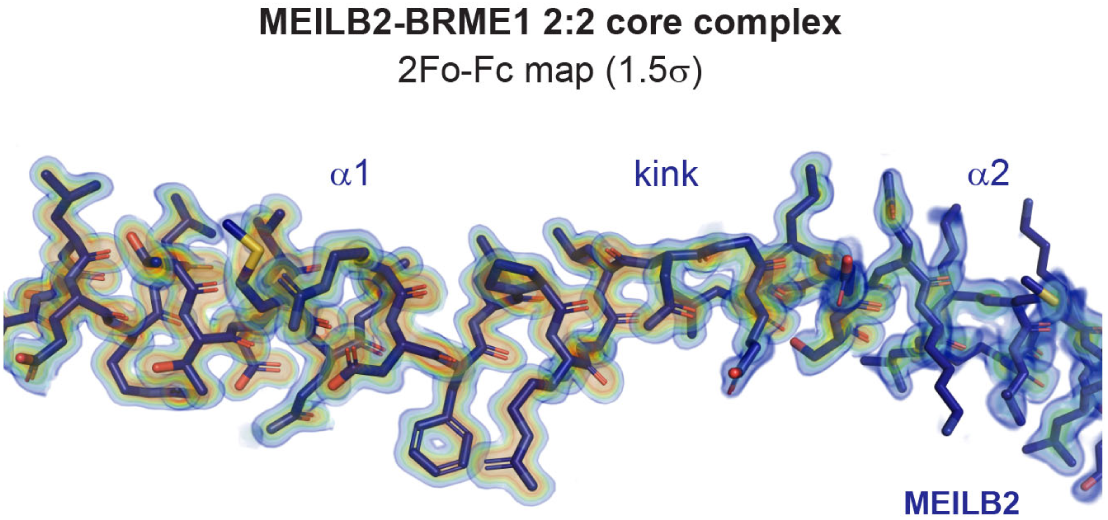
Crystal structure of the MEILB2-BRME1 2:2 core complex. 2Fo-Fc electron density map of the MEILB2-BRME1 2:2 core structure (1.5σ), presented as a rainbow between 1.5σ (blue) and 3.5σ (red), superimposed on the refined crystallographic model.

**Supplementary Figure 3.**
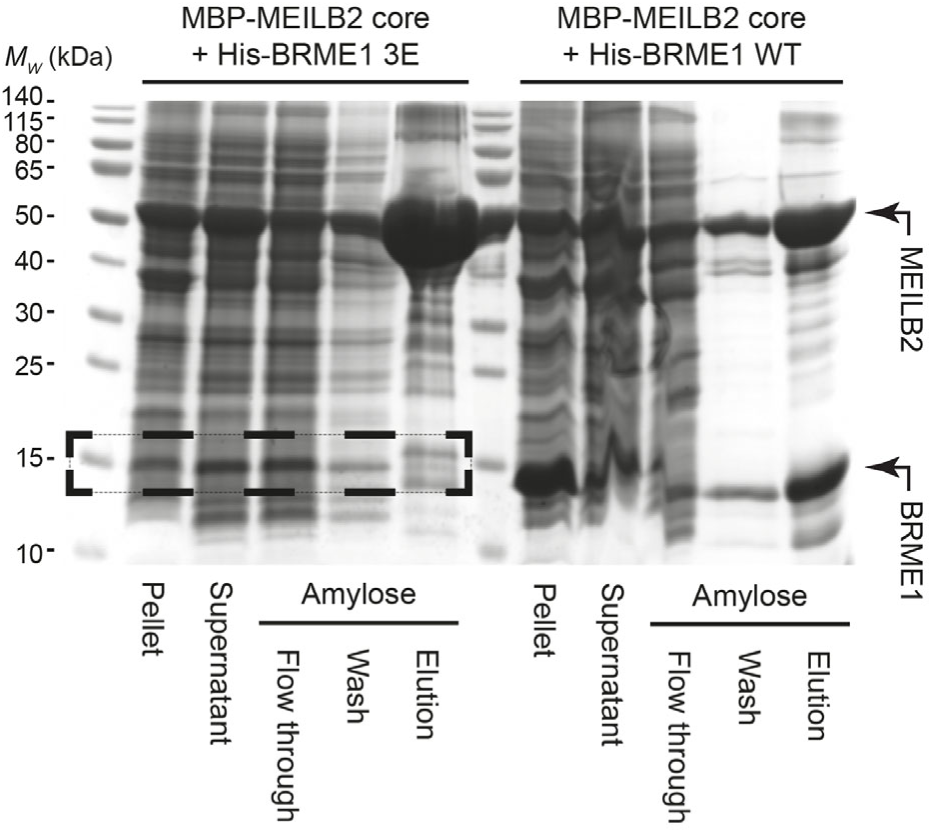
Pull-down of the MEILB2-BRME1 core complex. Amylose pull-down of His-BRME1 wild-type (WT) and His-BRME1 V548E L555E I562E (3E) with MBP-MEILB2 core following recombinant co-expression, corresponding to Figure 3b.

**Supplementary Figure 4.**
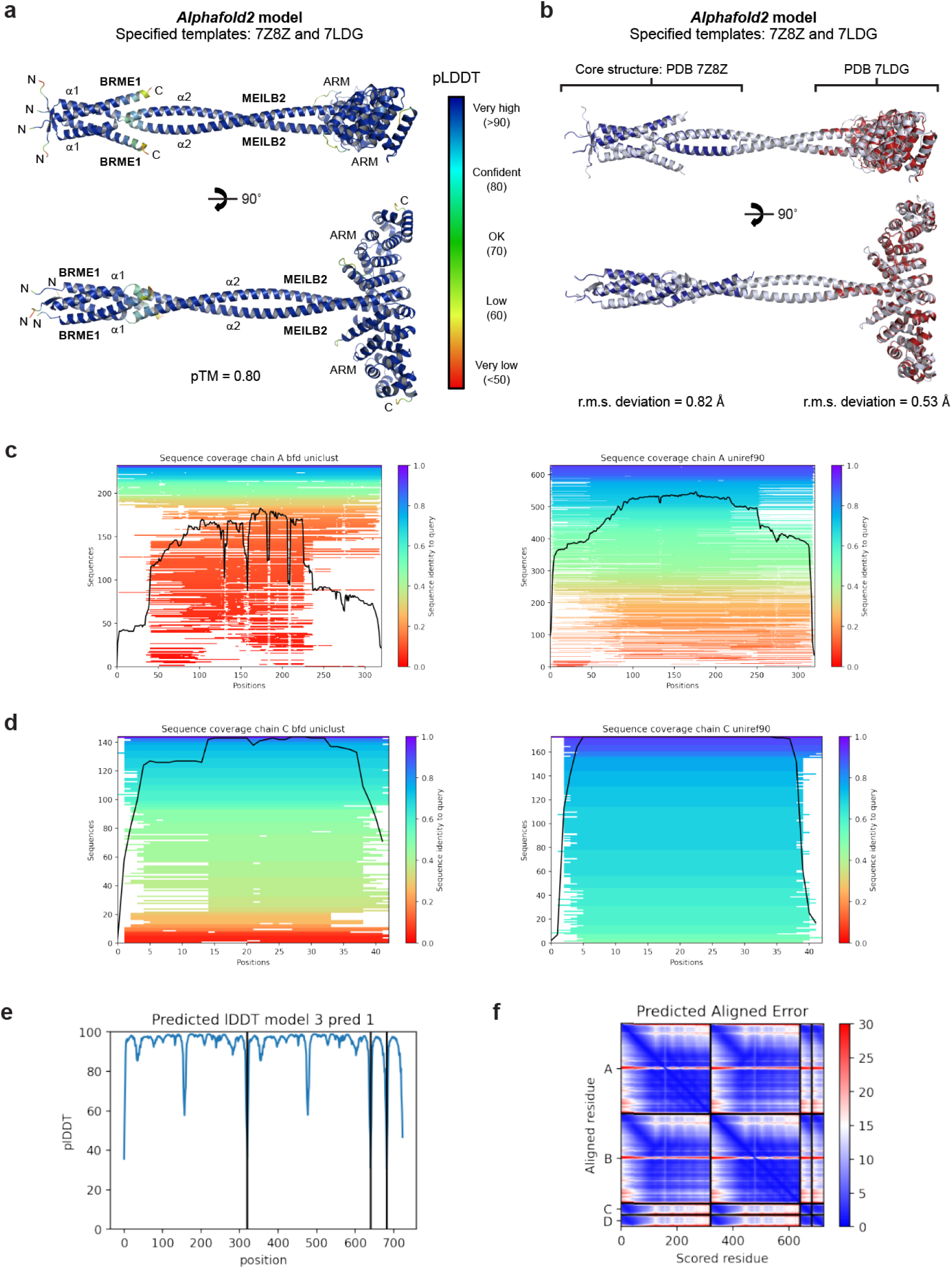
*Alphafold2* model of the full MEILB2-BRME1 2:2 complex. (**a**,**b**) *Alphafold2* model of the full MEILB2-BRME1 2:2 complex, generated using the MEILB2-BRME1 2:2 core structure reported herein (PDB accession 7Z8Z), and the MEILB2 ARM domain structure (PDB accession 7LDG; Pendlebury et al., 2021), as the sole templates. Corresponds to Figure 4a. (**a**) MEILB2-BRME1 2:2 model coloured according to predicted LDDT (pLDDT) scores, between blue (>90) and red (<50). (**b**) MEILB2-BRME1 2:2 model (light blue), with the superimposed MEILB2-BRME1 2:2 core crystal structure (dark blue; PDB accession 7Z8Z) and the MEILB2 ARM dimer crystal structure (red; PDB accession 7LDG) templates, showing r.m.s. deviations of 0.82 Å and 0.52 Å, respectively. (**c,d**) Representations of the multiple sequence alignments generated and used by *Alphafold2*, showing the number of sequences and sequence identity against the position along the (**c**) MEILB2 and (**d**) BRME1 query sequences. (**e**) Predicted LDDT (pLDDT) scores shown for each amino-acid of the two MEILB2 and two BRME1 chains. (**f**) Predicted aligned error scores between each amino-acid of the two MEILB2 and two BRME1 chains, between blue (low error) and red (high error).

**Supplementary Figure 5.**
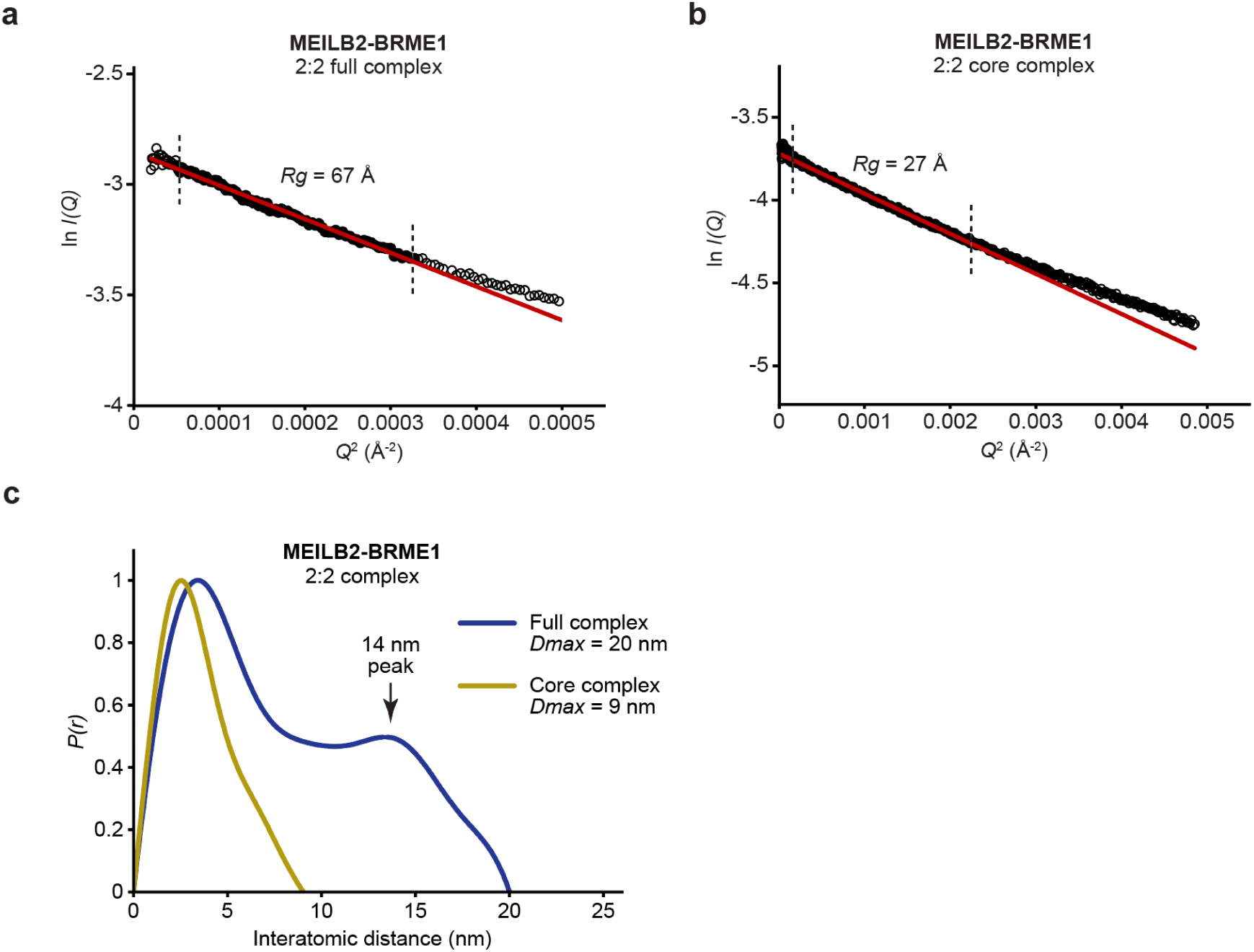
SEC-SAXS analysis of the MEILB2-BRME1 2:2 complex. (**a**,**b**) SAXS Guinier analysis of the MEILB2-BRME1 2:2 (**a**) full complex and (**b**) core complex to determine the radius of gyration (*Rg*); linear fits are shown in red, with the fitted data range highlighted in black and demarcated by dashed lines. The *Q*.*Rg* values were < 1.3 and *Rg* values were calculated as (**a**) 67 Å and (**b**) 27 Å. (**c**) SAXS *P(r)* interatomic distance distributions of the MEILB2-BRME1 2:2 full (blue) and core (yellow) complexes, showing maximum dimensions (*Dmax*) of 20 nm and 9 nm, respectively. The *P(r)* distribution for the full complex includes a peak at 14 nm, indicating that domains within the structure are separated by this distance.

**Supplementary Figure 6.**
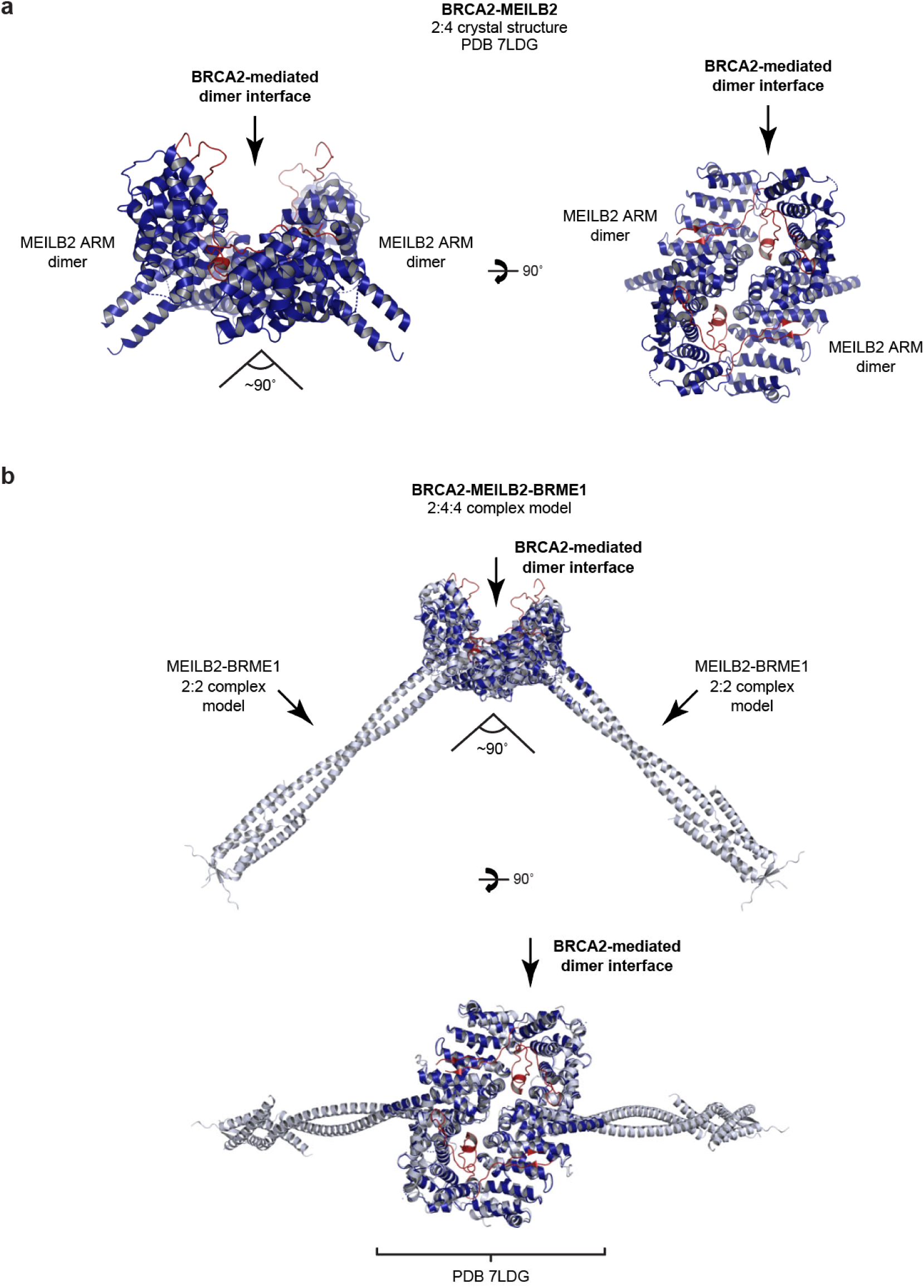
Model of the BRCA2-MEILB2-BRME1 2:4:4 ternary complex. (**a**) Crystal structure of the BRCA2-MEILB2 ARM 2:4 complex, showing how BRCA2 mediates the dimerization of two opposing MEILB2 ARM dimers, which held at approximately 90° to one another (PDB accession 7LDG; Pendlebury et al., 2021). (**b**) Model of the BRCA2-MEILB2-BRME1 2:4:4 ternary complex (corresponding to Figure 6b), generating by docking two models of the full MEILB2-BRME1 2:2 complex onto the BRCA2-MEILB2 ARM 2:4 crystal structure. In the resultant assembly, BRCA2 mediates dimerization of the two constituent MEILB2-BRME1 2:2 complex, in which their limbs are held at approximately 90° to one another.

**Supplementary Figure 7.**
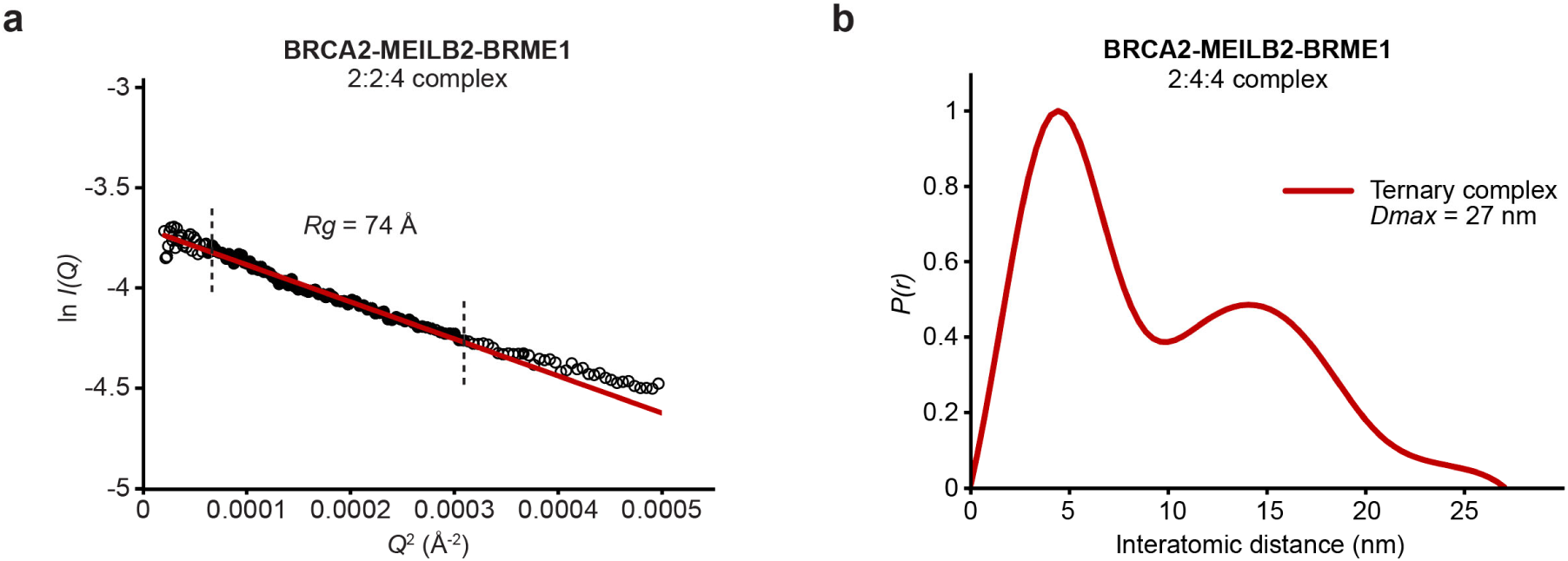
SEC-SAXS analysis of the BRCA2-MEILB2-BRME1 2:4:4 ternary complex. (**a**) SAXS Guinier analysis of the BRCA2-MEILB2-BRME1 2:4:4 ternary complex to determine the radius of gyration (*Rg*); the linear fit is shown in red, with the fitted data range highlighted in black and demarcated by dashed lines. The *Q*.*Rg* values were < 1.3 and *Rg* was calculated as 74 Å. (**c**) SAXS *P(r)* interatomic distance distribution of the BRCA2-MEILB2-BRME1 2:4:4 ternary complex, showing a maximum dimension (*Dmax*) of 27 nm.

## References

Adams, I. R. & Davies, O. R. 2023. Meiotic Chromosome Structure, the Synaptonemal Complex, and Infertility. Annu Rev Genomics Hum Genet.

Adams, P. D., Afonine, P. V., Bunkoczi, G., Chen, V. B., Davis, I. W., Echols, N., Headd, J. J., Hung, L. W., Kapral, G. J., Grosse-Kunstleve, R. W., Mccoy, A. J., Moriarty, N. W., Oeffner, R., Read, R. J., Richardson, D. C., Richardson, J. S., Terwilliger, T. C. & Zwart, P. H. 2010. Phenix: a comprehensive Python-based system for macromolecular structure solution. Acta Crystallogr D Biol Crystallogr, 66, 213–21.

Bell, J. C., Dombrowski, C. C., Plank, J. L., Jensen, R. B. & Kowalczykowski, S. C. 2023. BRCA2 chaperones RAD51 to single molecules of Rpa-coated ssdna. Proc Natl Acad Sci U S A, 120, e2221971120.

Brandsma, I., Sato, K., Van Rossum-Fikkert, S. E., Van Vliet, N., Sleddens, E., Reuter, M., Odijk, H., Van Den Tempel, N., Dekkers, D. H. W., Bezstarosti, K., Demmers, J. A. A., Maas, A., Lebbink, J., Wyman, C., Essers, J., Van Gent, D. C., Baarends, W. M., Knipscheer, P., Kanaar, R. & Zelensky, A. N. 2019. HSF2bp Interacts with a Conserved Domain of BRCA2 and Is Required for Mouse Spermatogenesis. Cell Rep, 27, 3790–3798 e7.

Caballero, I., Sammito, M., Millan, C., Lebedev, A., Soler, N. & Uson, I. 2018. Arcimboldo on coiled coils. Acta Crystallogr D Struct Biol, 74, 194–204.

Chen, V. B., Arendall, W. B., Headd, J. J., Keedy, D. A., Immormino, R. M., Kapral, G. J., Murray, L. W., Richardson, J. S. & Richardson, D. C. 2010. MolProbity: all-atom structure validation for macromolecular crystallography. Acta Crystallographica Section D-Biological Crystallography, 66, 12–21.

Dai, J., Voloshin, O., Potapova, S. & Camerini-Otero, R. D. 2017. Meiotic Knockdown and Complementation Reveals Essential Role of RAD51 in Mouse Spermatogenesis. Cell Rep, 18, 1383–1394.

Davies, O. R., Forment, J. V., Sun, M., Belotserkovskaya, R., Coates, J., Galanty, Y., Demir, M., Morton, C. R., Rzechorzek, N. J., Jackson, S. P. & Pellegrini, L. 2015. Ctip tetramer assembly is required for Dna-end resection and repair. Nat Struct Mol Biol, 22, 150–157.

Davies, O. R. & Pellegrini, L. 2007. Interaction with the BRCA2 C terminus protects RAD51-Dna filaments from disassembly by Brc repeats. Nat Struct Mol Biol, 14, 475–83.

Dragan, A. I. & Privalov, P. L. 2002. Unfolding of a leucine zipper is not a simple two-state transition. J Mol Biol, 321, 891–908.

Emsley, P., Lohkamp, B., Scott, W. G. & Cowtan, K. 2010. Features and development of Coot Acta Crystallogr D Biol Crystallogr, 66, 486–501.

Esashi, F., Galkin, V. E., Yu, X., Egelman, E. H. & West, S. C. 2007. Stabilization of RAD51 nucleoprotein filaments by the C-terminal region of BRCA2. Nat Struct Mol Biol, 14, 468–74.

Evans, P. R. 2011. An introduction to data reduction: space-group determination, scaling and intensity statistics. Acta Crystallogr D Biol Crystallogr, 67, 282–92.

Evans, R., O’neill, M., Pritzel, A., Antropova, N., Senior, A., Green, T., Žídek, A., Bates, R., Blackwell, S., Yim, J., Ronneberger, O., Bodenstein, S., Zielinski, M., Bridgland, A., Potapenko, A., Cowie, A., Tunyasuvunakool, K., Jain, R., Clancy, E., Kohli, P., Jumper, J. & Hassabis, D. 2021. Protein complex prediction with AlphaFold-Multimer. bioRxiv, 2021.10.04.463034.

Felipe-Medina, N., Caburet, S., Sanchez-Saez, F., Condezo, Y. B., De Rooij, D. G., Gomez, H. L., Garcia-Valiente, R., Todeschini, A. L., Duque, P., Sanchez-Martin, M. A., Shalev, S. A., Llano, E., Veitia, R. A. & Pendas, A. M. 2020. A missense in HSF2bp causing primary ovarian insufficiency affects meiotic recombination by its novel interactor C19ORF57/BRME1. Elife, 9.

Ghouil, R., Miron, S., Koornneef, L., Veerman, J., Paul, M. W., Le Du, M. H., Sleddens-Linkels, E., Van Rossum-Fikkert, S. E., Van Loon, Y., Felipe-Medina, N., Pendas, A. M., Maas, A., Essers, J., Legrand, P., Baarends, W. M., Kanaar, R., Zinn-Justin, S. & Zelensky, A. N. 2021. BRCA2 binding through a cryptic repeated motif to HSF2bp oligomers does not impact meiotic recombination. Nat Commun, 12, 4605.

Gnugge, R. & Symington, L. S. 2021. Dna end resection during homologous recombination. Curr Opin Genet Dev, 71, 99–105.

Hall, J. M., Lee, M. K., Newman, B., Morrow, J. E., Anderson, L. A., Huey, B. & King, M. C. 1990. Linkage of early-onset familial breast cancer to chromosome 17q21. Science, 250, 1684–9.

Hartmann, M. D., Mendler, C. T., Bassler, J., Karamichali, I., Ridderbusch, O., Lupas, A. N. & Hernandez Alvarez, B. 2016. alpha/beta coiled coils. Elife, 5.

Hunter, N. 2015. Meiotic Recombination: The Essence of Heredity. Cold Spring Harb Perspect Biol, 7.

Jensen, R. B., Carreira, A. & Kowalczykowski, S. C. 2010. Purified human BRCA2 stimulates RAD51-mediated recombination. Nature, 467, 678–83.

Jumper, J., Evans, R., Pritzel, A., Green, T., Figurnov, M., Ronneberger, O., Tunyasuvunakool, K., Bates, R., Zidek, A., Potapenko, A., Bridgland, A., Meyer, C., Kohl, S. A. A., Ballard, A. J., Cowie, A., Romera-Paredes, B., Nikolov, S., Jain, R., Adler, J., Back, T., Petersen, S., Reiman, D., Clancy, E., Zielinski, M., Steinegger, M., Pacholska, M., Berghammer, T., Bodenstein, S., Silver, D., Vinyals, O., Senior, A. W., Kavukcuoglu, K., Kohli, P. & Hassabis, D. 2021. Highly accurate protein structure prediction with AlphaFold. Nature, 596, 583–589.

Kabsch, W. 2010. Xds. Acta Crystallogr D Biol Crystallogr, 66, 125–32.

Keeney, S., Giroux, C. N. & Kleckner, N. 1997. Meiosis-specific Dna double-strand breaks are catalyzed by Spo11, a member of a widely conserved protein family. Cell, 88, 375–84.

Kowalczykowski, S. C. 2015. An Overview of the Molecular Mechanisms of Recombinational Dna Repair. Cold Spring Harb Perspect Biol, 7.

Lao, J. P. & Hunter, N. 2010. Trying to avoid your sister. PLos Biol, 8, e1000519.

Le, H. P., Ma, X., Vaquero, J., Brinkmeyer, M., Guo, F., Heyer, W. D. & Liu, J. 2020. DSS1 and ssdna regulate oligomerization of BRCA2. Nucleic Acids Res, 48, 7818–7833.

Li, M., Feng, H., Lin, Z., Zheng, J., Liu, D., Guo, R., Li, J., Li, R. H. W., Ng, E. H. Y., Huen, M. S. Y., Wang, P. J., Yeung, W. S. B. & Liu, K. 2020. The novel male meiosis recombination regulator coordinates the progression of meiosis prophase I. J Genet Genomics, 47, 451–465.

Liu, J., Doty, T., Gibson, B. & Heyer, W. D. 2010. Human BRCA2 protein promotes RAD51 filament formation on Rpa-covered single-stranded Dna. Nat Struct Mol Biol, 17, 1260–2.

Luo, M., Yang, F., Leu, N. A., Landaiche, J., Handel, M. A., Benavente, R., La Salle, S. & Wang, P. J. 2013. Meiob exhibits single-stranded Dna-binding and exonuclease activities and is essential for meiotic recombination. Nat Commun, 4, 2788.

Madeira, F., Park, Y. M., Lee, J., Buso, N., Gur, T., Madhusoodanan, N., Basutkar, P., Tivey, A. R. N., Potter, S. C., Finn, R. D. & Lopez, R. 2019. The Embl-Ebi search and sequence analysis tools APIs in 2019. Nucleic Acids Res, 47, W636–W641.

Marston, N. J., Richards, W. J., Hughes, D., Bertwistle, D., Marshall, C. J. & Ashworth, A.1999. Interaction between the product of the breast cancer susceptibility gene BRCA2 and DSS1, a protein functionally conserved from yeast to mammals. Mol Cell Biol, 19, 4633–42.

Martinez, J. S., Von Nicolai, C., Kim, T., Ehlen, A., Mazin, A. V., Kowalczykowski, S. C. & Carreira, A. 2016. BRCA2 regulates DMC1-mediated recombination through the Brc repeats. Proc Natl Acad Sci U S A, 113, 3515–20.

Mccoy, A. J., Grosse-Kunstleve, R. W., Adams, P. D., Winn, M. D., Storoni, L. C. & Read, R. J.2007. Phaser crystallographic software. Journal of Applied Crystallography, 40, 658–674.

Mirdita, M., Schutze, K., Moriwaki, Y., Heo, L., Ovchinnikov, S. & Steinegger, M. 2022. ColabFold: making protein folding accessible to all. Nat Methods, 19, 679–682.

Nasmyth, K. 2005. How might cohesin hold sister chromatids together? Philos Trans R Soc Lond B Biol Sci, 360, 483–96.

Oh, J. & Symington, L. S. 2018. Role of the Mre11 Complex in Preserving Genome Integrity. Genes (Basel), 9.

P.V.Konarev, V. V. V., A.V. Sokolova, M.H.J. Koch, D. I. Svergun 2003. Primus - a Windows-Pc based system for small-angle scattering data analysis. J Appl Cryst., 36, 1277–1282

Patel, K. J., Yu, V. P., Lee, H., Corcoran, A., Thistlethwaite, F. C., Evans, M. J., Colledge, W. H., Friedman, L. S., Ponder, B. A. & Venkitaraman, A. R. 1998. Involvement of Brca2 in Dna repair. Mol Cell, 1, 347–57.

Paul, M. W., Sidhu, A., Liang, Y., Van Rossum-Fikkert, S. E., Odijk, H., Zelensky, A. N., Kanaar, R. & Wyman, C. 2021. Role of BRCA2 Dna-binding and C-terminal domain in its mobility and conformation in Dna repair. Elife, 10.

Pellegrini, L., Yu, D. S., Lo, T., Anand, S., Lee, M., Blundell, T. L. & Venkitaraman, A. R. 2002. Insights into Dna recombination from the structure of a RAD51-BRCA2 complex. Nature, 420, 287–93.

Pendlebury, D. F., Zhang, J., Agrawal, R., Shibuya, H. & Nandakumar, J. 2021. Structure of a meiosis-specific complex central to BRCA2 localization at recombination sites. Nat Struct Mol Biol, 28, 671–680.

Peranen, J., Rikkonen, M., Hyvonen, M. & Kaariainen, L. 1996. T7 vectors with modified T7lac promoter for expression of proteins in Escherichia coli. Anal Biochem, 236, 371–3.

Piazza, A., Bordelet, H., Dumont, A., Thierry, A., Savocco, J., Girard, F. & Koszul, R. 2021. Cohesin regulates homology search during recombinational Dna repair. Nat Cell Biol, 23, 1176–1186.

Pittman, D. L., Cobb, J., Schimenti, K. J., Wilson, L. A., Cooper, D. M., Brignull, E., Handel, M. A. & Schimenti, J. C. 1998. Meiotic prophase arrest with failure of chromosome synapsis in mice deficient for Dmc1, a germline-specific Reca homolog. Mol Cell, 1, 697–705.

Ribeiro, J., Dupaigne, P., Petrillo, C., Ducrot, C., Duquenne, C., Veaute, X., Saintome, C., Busso, D., Guerois, R., Martini, E. & Livera, G. 2021. The meiosis-specific Meiob- SPATA22 complex cooperates with Rpa to form a compacted mixed Meiob/SPATA22/Rpa/ssdna complex. Dna Repair (Amst), 102, 103097.

Robert, T., Nore, A., Brun, C., Maffre, C., Crimi, B., Bourbon, H. M. & De Massy, B. 2016. The Topovib-Like protein family is required for meiotic Dna double-strand break formation. Science, 351, 943–9.

Rodriguez, D. D., Grosse, C., Himmel, S., Gonzalez, C., De Ilarduya, I. M., Becker, S., Sheldrick, G. M. & Uson, I. 2009. Crystallographic ab initio protein structure solution below atomic resolution. Nature Methods, 6, 651–U39.

Sato, K., Brandsma, I., Van Rossum-Fikkert, S. E., Verkaik, N., Oostra, A. B., Dorsman, J. C., Van Gent, D. C., Knipscheer, P., Kanaar, R. & Zelensky, A. N. 2020. HSF2bp negatively regulates homologous recombination in Dna interstrand crosslink repair. Nucleic Acids Res, 48, 2442–2456.

Shahid, T., Soroka, J., Kong, E., Malivert, L., Mcilwraith, M. J., Pape, T., West, S. C. & Zhang, X. 2014. Structure and mechanism of action of the BRCA2 breast cancer tumor suppressor. Nat Struct Mol Biol, 21, 962–968.

Shang, Y., Huang, J., Li, W., Zhang, Y., Zhou, X., Shao, Q., Tan, T., Yin, S., Zhang, L. & Wang, S. 2022. MEIOK21 regulates oocyte quantity and quality via modulating meiotic recombination. Faseb J, 36, e22357.

Shang, Y., Huang, T., Liu, H., Liu, Y., Liang, H., Yu, X., Li, M., Zhai, B., Yang, X., Wei, Y., Wang, G., Chen, Z., Wang, S. & Zhang, L. 2020. MEIOK21: a new component of meiotic recombination bridges required for spermatogenesis. Nucleic Acids Res, 48, 6624–6639.

Sharan, S. K., Pyle, A., Coppola, V., Babus, J., Swaminathan, S., Benedict, J., Swing, D., Martin, B. K., Tessarollo, L., Evans, J. P., Flaws, J. A. & Handel, M. A. 2004. BRCA2 deficiency in mice leads to meiotic impairment and infertility. Development, 131, 131–42.

Shibuya, H., Morimoto, A. & Watanabe, Y. 2014. The dissection of meiotic chromosome movement in mice using an in vivo electroporation technique. PLos Genet, 10, e1004821.

Shinohara, A. & Shinohara, M. 2004. Roles of Reca homologues Rad51 and Dmc1 during meiotic recombination. Cytogenet Genome Res, 107, 201–7.

Souquet, B., Abby, E., Herve, R., Finsterbusch, F., Tourpin, S., Le Bouffant, R., Duquenne, C., Messiaen, S., Martini, E., Bernardino-Sgherri, J., Toth, A., Habert, R. & Livera, G. 2013. Meiob targets single-strand Dna and is necessary for meiotic recombination. PLos Genet, 9, e1003784.

Svergun D.I. B. C. K. M. H. J. 1995. Crysol – a Program to Evaluate X-ray Solution Scattering of Biological Macromolecules from Atomic Coordinates. J. Appl. Cryst., 28, 768–773.

Takemoto, K., Tani, N., Takada-Horisawa, Y., Fujimura, S., Tanno, N., Yamane, M., Okamura, K., Sugimoto, M., Araki, K. & Ishiguro, K. I. 2020. Meiosis-Specific C19orf57/4930432K21Rik/BRME1 Modulates Localization of RAD51 and DMC1 to DSBs in Mouse Meiotic Recombination. Cell Rep, 31, 107686.

Thorslund, T., Esashi, F. & West, S. C. 2007. Interactions between human BRCA2 protein and the meiosis-specific recombinase DMC1. Embo J, 26, 2915–22.

Thorslund, T., Mcilwraith, M. J., Compton, S. A., Lekomtsev, S., Petronczki, M., Griffith, J. D. & West, S.C.. 2010. The breast cancer tumor suppressor BRCA2 promotes the specific targeting of RAD51 to single-stranded Dna. Nat Struct Mol Biol, 17, 1263–5.

Vonrhein, C., Flensburg, C., Keller, P., Sharff, A., Smart, O., Paciorek, W., Womack, T. & Bricogne, G. 2011. Data processing and analysis with the autoproc toolbox. Acta Crystallogr D Biol Crystallogr, 67, 293–302.

Waterhouse, A. M., Procter, J. B., Martin, D. M., Clamp, M. & Barton, G. J. 2009. Jalview Version 2--a multiple sequence alignment editor and analysis workbench. Bioinformatics, 25, 1189–91.

Wilkinson, O. J., Martin-Gonzalez, A., Kang, H., Northall, S. J., Wigley, D. B., Moreno-Herrero, F. & Dillingham, M. S. 2019. Ctip forms a tetrameric dumbbell-shaped particle which bridges complex Dna end structures for double-strand break repair. Elife, 8.

Yang, H., Jeffrey, P. D., Miller, J., Kinnucan, E., Sun, Y., Thoma, N. H., Zheng, N., Chen, P. L., Lee, W. H. & Pavletich, N. P. 2002. BRCA2 function in Dna binding and recombination from a BRCA2-DSS1-ssdna structure. Science, 297, 1837–48.

Yu, V. P., Koehler, M., Steinlein, C., Schmid, M., Hanakahi, L. A., Van Gool, A. J., West, S. C. & Venkitaraman, A. R. 2000. Gross chromosomal rearrangements and genetic exchange between nonhomologous chromosomes following BRCA2 inactivation. Genes Dev, 14, 1400–6.

Zhang, J., Fujiwara, Y., Yamamoto, S. & Shibuya, H. 2019. A meiosis-specific BRCA2 binding protein recruits recombinases to Dna double-strand breaks to ensure homologous recombination. Nat Commun, 10, 722.

Zhang, J., Gurusaran, M., Fujiwara, Y., Zhang, K., Echbarthi, M., Vorontsov, E., Guo, R., Pendlebury, D. F., Alam, I., Livera, G., Emmanuelle, M., Wang, P. J., Nandakumar, J., Davies, O. R. & Shibuya, H. 2020. The BRCA2-MEILB2-BRME1 complex governs meiotic recombination and impairs the mitotic BRCA2-RAD51 function in cancer cells. Nat Commun, 11, 2055.

Zhang, J., Nandakumar, J. & Shibuya, H. 2022. BRCA2 in mammalian meiosis. Trends Cell Biol, 32, 281–284.

Zhao, W., Vaithiyalingam, S., San Filippo, J., Maranon, D. G., Jimenez-Sainz, J., Fontenay, G. V., Kwon, Y., Leung, S. G., Lu, L., Jensen, R. B., Chazin, W. J., Wiese, C. & Sung, P. 2015.Promotion of BRCA2-Dependent Homologous Recombination by DSS1 via Rpa Targeting and Dna Mimicry. Mol Cell, 59, 176–87.

Zickler, D. & Kleckner, N. 2015. Recombination, Pairing, and Synapsis of Homologs during Meiosis. Cold Spring Harb Perspect Biol, 7.

